# Rapid development of SARS-CoV-2 receptor binding domain-conjugated nanoparticle vaccine candidate

**DOI:** 10.1101/2020.11.03.366138

**Authors:** Yin-Feng Kang, Cong Sun, Zhen Zhuang, Run-Yu Yuan, Qing-Bing Zheng, Jiang-Ping Li, Ping-Ping Zhou, Xin-Chun Chen, Xiao Zhang, Xiao-Hui Yu, Xiang-Wei Kong, Qian-Ying Zhu, Miao Xu, Nan-Shan Zhong, Yi-Xin Zeng, Guo-Kai Feng, Chang-Wen Ke, Jin-Cun Zhao, Mu-Sheng Zeng

**Affiliations:** State Key Laboratory of Oncology in South China, Collaborative Innovation Center for Cancer Medicine, Guangdong Key Laboratory of Nasopharyngeal Carcinoma Diagnosis and Therapy, Department of Experimental Research, Sun Yat-sen University Cancer Center, Sun Yat-sen University, Guangzhou, Guangdong, P. R. China; State Key Laboratory of Respiratory Disease, National Clinical Research Center for Respiratory Disease, Guangzhou Institute of Respiratory Health, the First Affiliated Hospital of Guangzhou Medical University, Guangzhou, P. R. China; Guangdong Provincial Institution of Public Health, Guangdong Provincial Center for Disease Control and Prevention, Guangzhou, Guangdong, P. R. China; State Key Laboratory of Molecular Vaccinology and Molecular Diagnostics, National Institute of Diagnostics and Vaccine Development in Infectious Diseases, School of Public Health, Xiamen University, Xiamen, Fujian, PR China

**Keywords:** SARS-CoV-2, RBD, Spytag-SpyCatcher, nanoparticle

## Abstract

The ongoing of coronavirus disease 2019 (COVID-19) pandemic caused by novel SARS-CoV-2 coronavirus, resulting in economic losses and seriously threating the human health in worldwide, highlighting the urgent need of a stabilized, easily produced and effective preventive vaccine. The SARS-COV-2 spike protein receptor binding region (RBD) plays an important role in the process of viral binding receptor angiotensin-converting enzyme 2 (ACE2) and membrane fusion, making it an ideal target for vaccine development. In this study, we designed three different RBD-conjugated nanoparticles vaccine candidates, RBD-Ferritin (24-mer), RBD-mi3 (60-mer) and RBD-I53-50 (120-mer), with the application of covalent bond linking by SpyTag-SpyCatcher system. It was demonstrated that the neutralizing capability of sera from mice immunized with three RBD-conjugated nanoparticles adjuvanted with AddaVax or Sigma Systerm Adjuvant (SAS) after each immunization was ~8-to 120-fold greater than monomeric RBD group in SARS-CoV-2 pseudovirus and authentic virus neutralization assay. Most importantly, sera from RBD-conjugated NPs groups more efficiently blocked the binding of RBD to ACE2 or neutralizing antibody in vitro, a further proof of promising immunization effect. Besides, high physical stability and flexibility in assembly consolidated the benefit for rapid scale-up production of vaccine. These results supported that our designed SARS-CoV-2 RBD-conjugated nanoparticle was competitive vaccine candidate and the carrier nanoparticles could be adopted as universal platform for future vaccine development.

## 1. Introduction

The unexpected outbreak of COVID-19 pandemic since 2019 has become a global public health crisis affecting 216 countries or regions, 26016839 confirmed cases and over 850000 confirmed death (https://www.who.int/emergencies/diseases/novel-coronavirus-2019). SARS-CoV-2, the causative virus affirmed by laboratory evaluation and comprehensive sequencing, belongs to β-coronavirus in coronavirus family which comprises other highly-pathogenetic virus strains (SARS-CoV, MERS-CoV) for humans ^1–3^. The infection caused a diverse clinical characterization of respiratory syndrome and person-to-person transmission ^4^, and even led to death ^5–6^.

As a member of coronavirus, SARS-CoV-2 adopts a similar cell-entry mechanism relying on spike protein on viral membrane to fulfill the host cell recognition, attachment and membrane fusion, while the spike protein of coronavirus also shared great similarity in structural appearance as a trimeric fusion protein ^7–9^. Specifically, SARS-CoV-2 spike protein recognize the angiotensin converting enzyme 2 (ACE2) as the entry receptor like SARS-CoV and the key binding interface lies on the receptor binding domain (RBD) of the spike protein, which has been confirmed by both structural elucidation through high resolution Cryo-EM structure and interface mutation scanning in previous work ^10–11^.

With the basis of clear structural information and biological function of SARS-CoV-2 spike protein and the key region RBD, most neutralizing antibodies and potential therapeutic agent are found against the spike protein RBD, making this region an ideal target for vaccine development ^8, 12–13^. However, despite a comprehensive effort on RBD-based vaccine, application of RBD subunit is still hindered by the quite low immunogenicity due to a variety of reason ^14^. To increase the immunogenicity, scientists endeavor to modify the RBD to achieve larger antigen-carrier complex size or multimerization, which would complicate the overall structure of RBD and prolong the production and validation process of the recombinant modified antigen ^15–16^. To shorten the time cost of vaccine development during emergency, a more compact workflow of antigen-displayed nanoparticle production is in demand. Protein covalent bond linking strategy has enjoyed a rapid development in recent years, rendering greater easiness to protein modification and multimerization ^17–19^. SpyTag-SpyCatcher system has been used as a strategy for antigen display on particles in hepatitis B ^20^ and HIV ^21^ vaccine development, overcoming the obstacles in massive in vitro production of fusion protein of antigen and nanoparticle scaffold.

Here we reported our designed SARS-CoV-2 spike protein RBD-based nanoparticle vaccines in application of the covalent bond linking strategy. With SpyTag fused to the C-termius of RBD, the antigen could covalently be linked to the SpyCatcher upon the nanoparticle scaffold. The RBD nanoparticles could elicit higher neutralizing antibody titers compared to monomer RBD in mice, confirmed by stronger sera RBD-competition with both ACE2 and neutralizing antibody. Besides, our work validated 3 different nanoparticles platform with SpyCatcher in N-terminus viable for SpyTag-fused protein coupling, which could become general nanoparticle capture platform for other antigen in the future.

## 2. Results

### 2.1. Design and production of RBD-conjugated nanoparticles

Previous studies have demonstrated that immunization with receptor binding region (RBD) of SARS-CoV-2 Spike protein formulated with aluminum hydroxide adjuvant in mice elicited higher neutralization antibody titers in comparison with the extracellular domain protein (ECD), S1-subunit protein (S1) and S2-subunit protein (S2) ^22^. In addition, RBD amino acid sequences from 24 representative SARS CoV-2 strains isolated in different countries were aligned and founded to be very conservative (Figure S1, Supporting Information). In the present study, we focused on the RBD of SARS-CoV-2 S glycoprotein to design the RBD-conjugated nanoparticle vaccine based on the SpyTag-SpyCatcher system. As mentioned in previous study^17–18^, the enhanced shortening form of SpyCatcher, ΔN1-SpyCatcher was used in our study. Therefore, we engineered and adapted three ΔN1-SpyCatcher-nanoparticles (Δ N1-SpyCatcher-NPs) conjugation platform, Δ N1-SpyCatcher-Ferritin, ΔN1-SpyCatcher-mi3 and ΔN1-SpyCatcher-I53-50, to display more antigen to the surface of NPs based on the formation of isopeptide bond between SpyTag peptide and ΔN1-SpyCatcher in vivo (Figure S2b, Supporting Information). 24-mers ferritin were self-assembled into a spherical particle and form an octahedral nanocage ^23^. Computational designed and optimized mi3 NP protein with mutation of C76A and C100A from KDPG aldolases to escape the potential disulfide bond-mediated heterogeneity was dodecameric cage engineered scaffold with 60 total subunits multiply display its on surface of NPs ^17, 24^. The I53-50 NPs was a computational designed icosahedral nanoparticle assembled with two components, 20 copies of trimeric I53-50A1.1PT1 and 12 copiesof pentameric I53-50B.4PT1 ^25^.

Previous studies have manifested that immunization with RBD of SARS-CoV S protein expressed in the mammalian cells could elicited higher potent neutralizing antibody responses in mice and provided completely protection following infection with SARS-CoV compared with those expressed in insect cells and E.coli ^26^. RBD-SpyTag (residues 319–541) protein was firstly expressed by transient transfection method into HEK293F cells and purified by Ni-NTA affinity chromatography, and followed by SEC. As shown in Figure 1C and 1D, purified RBD-SpyTag was identified uniform and highly pure demonstrated by a clear single blot in SDS-PAGE and a single major peak in SEC chromatogram. As shown in Figure 1C and 1D, ΔN1-SpyCatcher-NPs, ΔN1-SpyCatcher-Ferritin NP, ΔN1-SpyCatcher-mi3 NP and Δ N1-SpyCatcher-I53-50 NP were expressed in E.coli and purified by Ni-NTA affinity chromatography followed by SEC, and the high yield and production quality of protein could be observed also from the SDS-PAGE and SEC peak result. Further, with the preparation of precursor proteins in high quality produced, assembly of the RBD-conjugated nanoparticles, purified Δ N1-SpyCatcher-Ferritin, ΔN1-SpyCatcher-mi3 and ΔN1-SpyCatcher-I53-50A1.1PT1 proteins was performed. A 50 μM subunit concentration of RBD-SpyTag was incubated with 8 times higher excess molar of ΔN1-SpyCatcher-NPs overnight for in vitro bonding reaction, and then applied to SEC assay to separate out RBD monomers and unlinked nanoparticles. As shown in Figure. 1C RBD-conjugated Ferritin, mi3 and I53-50 NPs were verified through the single band and an uniformly increase of molecular weight (around 35 kDa to 72 kDa) from reducing SDS-PAGE and further confirmed by peak forward shifts from the SEC assay (Figure 1D), which suggested that RBD-SpyTag could completely conjugated with ΔN1-SpyCatcher-NPs at a high efficacy and full reaction level. Collectively, we used the SpyTag-SpyCatcher system to guarantee a both flexible and high-efficiency production of SARS-CoV-2 RBD-conjugated nanoparticles.

**Figure 1.**
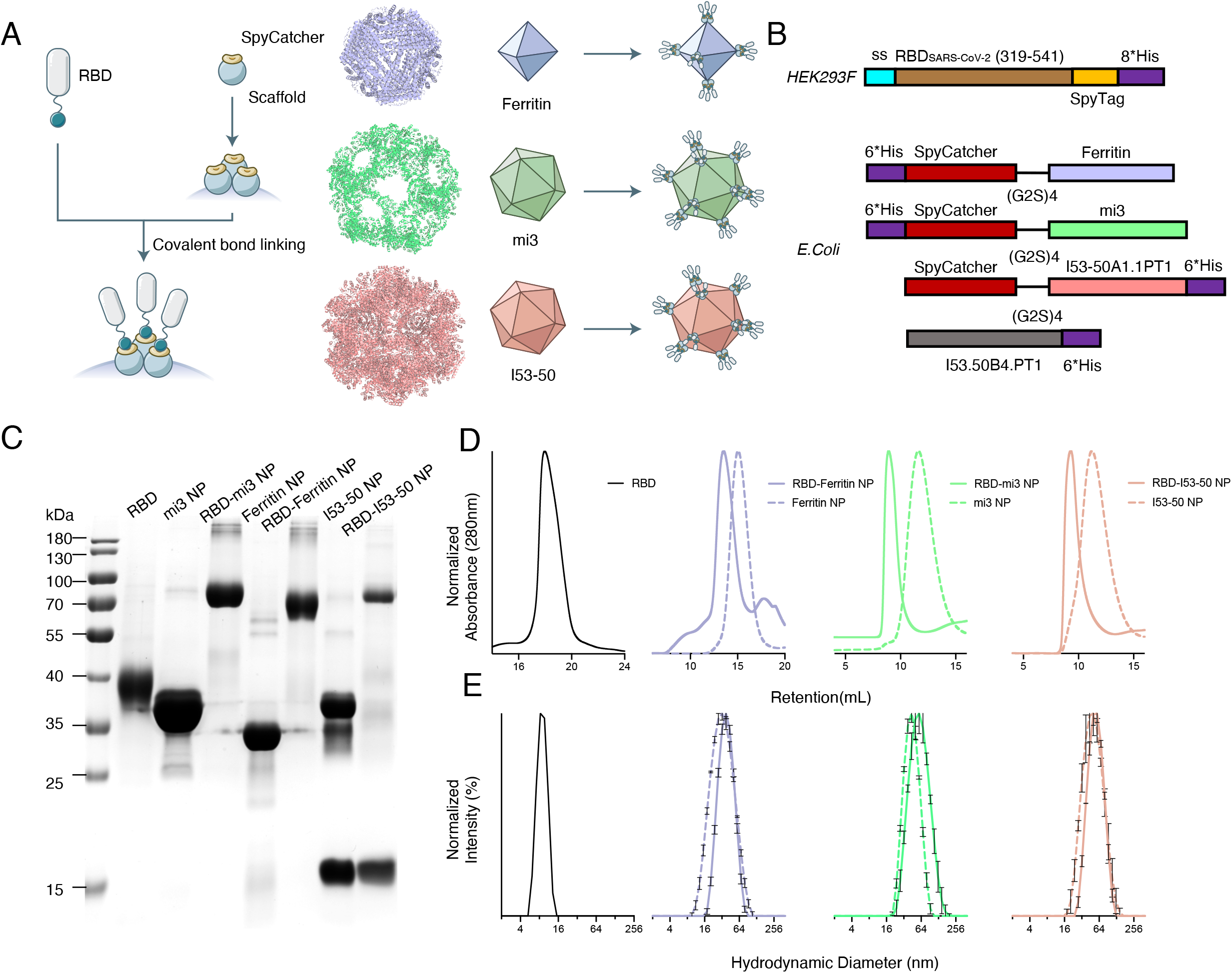
Construction and structural characteristics of RBD-conjugated nanoparticles. (A) Sketch of RBD nanoparticle design. The left flow diagram shows a brief introduction to the modification to RBD and nanoparticle scaffolds with fusion of SpyTag-SpyCatcher system. The right schema display ideal nanoparticles with full valency of RBD. Colors of each nanoparticle is accordant to the displayed palette of the following charts. (B) Construction of target protein expression plasmid in different expression system,E. coli and HEK293F. (C) Reduced SDS-PAGE of the RBD monomer, RBD-conjugated NPs and unbonded nanoparticles. A high covalent bond linking efficiency is achieved as the blot of RBD monomer and unlinked nanoparticle scaffold disappear in the lane of RBD-conjugated NPs. (D) Size exclusion chromatography (SEC) of RBD monomer, RBD-conjugated NPs and unbonded nanoparticles on Superose 6 increase 10/300GL. Peak forward shifts of retention are observed after bond linking of RBD-SpyTag and Δ N1-SpyCatcher-NPs. (E) Dynamic light scattering (DLS) of RBD monomer, RBD-conjugated NPs and unbonded nanoparticles. Increased hydrodynamics diameters of nanoparticles after bond linking are shown.

### 2.2. Structural characterization of SARS-CoV-2 RBD-conjugated nanoparticles

We next observe the structural characterization of RBD-conjugated nanoparticles by using negative stain electron microscopic (EM). As shown in Figure 2A, RBD was conjugated with Ferritin, mi3 and I53-50 nanoparticles and presented on the surface of monodispersed particles. As shown by the EM graphs, a burred exterior surface of nanoparticles could be observed among RBD-conjugated NPs, especially for RBD-Ferritin NP. The hydrodynamic diameters of RBD monomer, ΔN1-SpyCatcher-NPs and RBD-conjugated NPs were further measured by Dynamic Light Scattering (DLS). As displayed in Figure 2B, particle characteristic of both unconjugated NPs and RBD-conjugated NPs were validated and uniform distribution of particle sizes were rendered. Moreover, consistent with the results of negative stain EM, the hydrodynamic diameter of RBD-conjugated nanoparticles was larger than the unconjugated NPs from DLS.

**Figure 2.**
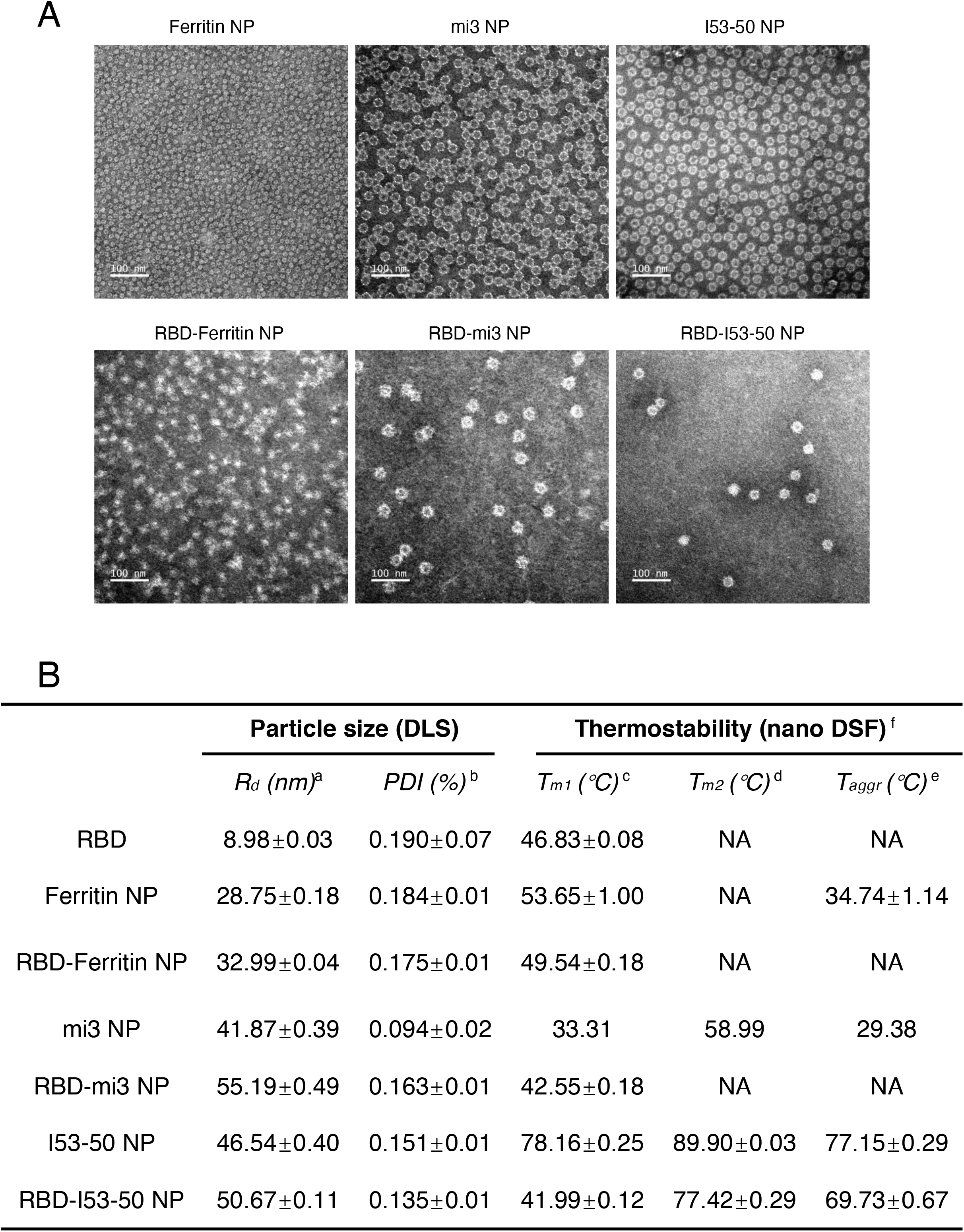
Assembly validation and physical evaluation of nanoparticles. (A) Negative stain electron micrographs of unlinked nanoparticles and RBD-conjugated NPs. (B) Detailed information of DLS and nano DSF results. a. R_d_: Hydrodynamics diameter b. PDI: Polydispersity index, PDI lower than 0.2 indicates a uniform particle size. c. T_m1_: the first melting temperature d. T_m2_: the second melting temperature e. T_aggr_: the aggregation temperature f. Melting temperature and aggregation temperature are given by the analysis software of nanoDSF

In order to explore the physical stability of the nanoparticles to verify the compatibility of antigen and the nanoparticle platform, nano differential scanning fluorimetry (nanoDSF) was performed to RBD monomer, Δ N1-SpyCatcher-NPs and RBD-conjugated NPs, and detailed thermostability parameters were given (Figure 2B). A close Tm1 of RBD and RBD-conjugated NPs primarily indicated that the overall structure of RBD upon the nanoparticle was not affected by the conjugation. The risen Tm1 for mi3-NP after conjugation with RBD may be ascribed to RBD-buried unsatisfied exterior surface, which even strengthened the structural viability of covalent bond linking strategy for RBD with the nanoparticles. Except for melting temperatures, no aggregation was observed for RBD monomer, RBD-Ferritin NP and RBD-mi3 NP during the process of thermal denaturation (Figure 2B). Comparatively, RBD-I53-50 NP underwent aggregation under approximately 70°C, close to the aggregation temperature of empty I53-50 NP and significantly higher than the Tm1 of the RBD monomer. The above results demonstrated that under general medicine or vaccine storage environment at 4°C, the designed conjugated vaccine could maintain a similar stability behavior as the RBD monomer which highly benefited commercial production and distribution.

### 2.3. In vitro antigenicity validation of SARS-CoV-2 RBD-conjugated nanoparticles

We next expressed and purified recombinant human ACE2 ectodomain and RBD-specific neutralizing antibody (CB6), and then characterized the antigenicity of RBD-conjugated NPs by detecting the binding affinity with the receptor and antibody. CB6 neutralization antibody was isolated from a COVID-19 convalescent patient and recognized an epitope that overlap with the hACE2-binding site of RBD, a critical character enabling a potential effect to neutralize the authentic SARS-CoV-2 virus ^27^. ELISA profiles showed that RBD-SpyTag monomer and three RBD-conjugated NPs bound to hACE2 and CB6 antibody in a dose-dependent manner. Analogous to soluble RBD-SpyTag monomer, three RBD-conjugated NPs bound to purified hACE2, suggesting that the conformation of RBD monomer was retained on the conjugated nanoparticles (Figure 3A). However, the binding between three RBD-conjugated NPs and CB6 antibody was significantly higher than RBD-SpyTag monomer (Figure 3B). Bio-layer interferometry assay was then applied to further examine the binding kinetics of RBD-conjugated NPs. As illustrated in Figure 3B and 3D, the measured binding affinity constant (kD) of RBD monomer and two RBD-conjugated NPs, RBD-Ferritin NPand RBD-I53-50 NP, with hACE2 receptor were 4.34E-09, 1.74E-08 and 1.00E-09 M. However, the dissociation was very slowly between RBD-mi3 NP and hACE2, and kD reached up to 1.0E-12 M, indicating RBD-mi3 NP showed an even higher antigenicity in comparison to RBD-Ferritin NP and RBD-I53-50 NP. The binding kinetics of RBD-conjugated NPs to CB6 antibody were also measured. The binding capability between three RBD-conjugated NPs and CB6 antibody were significantly stronger than RBD monomer (Figure 3C and 3D), suggesting that three RBD-conjugated NPs may showed a higher affinity to specific BCR targeting the RBD of SARS-CoV-2.

**Figure 3.**
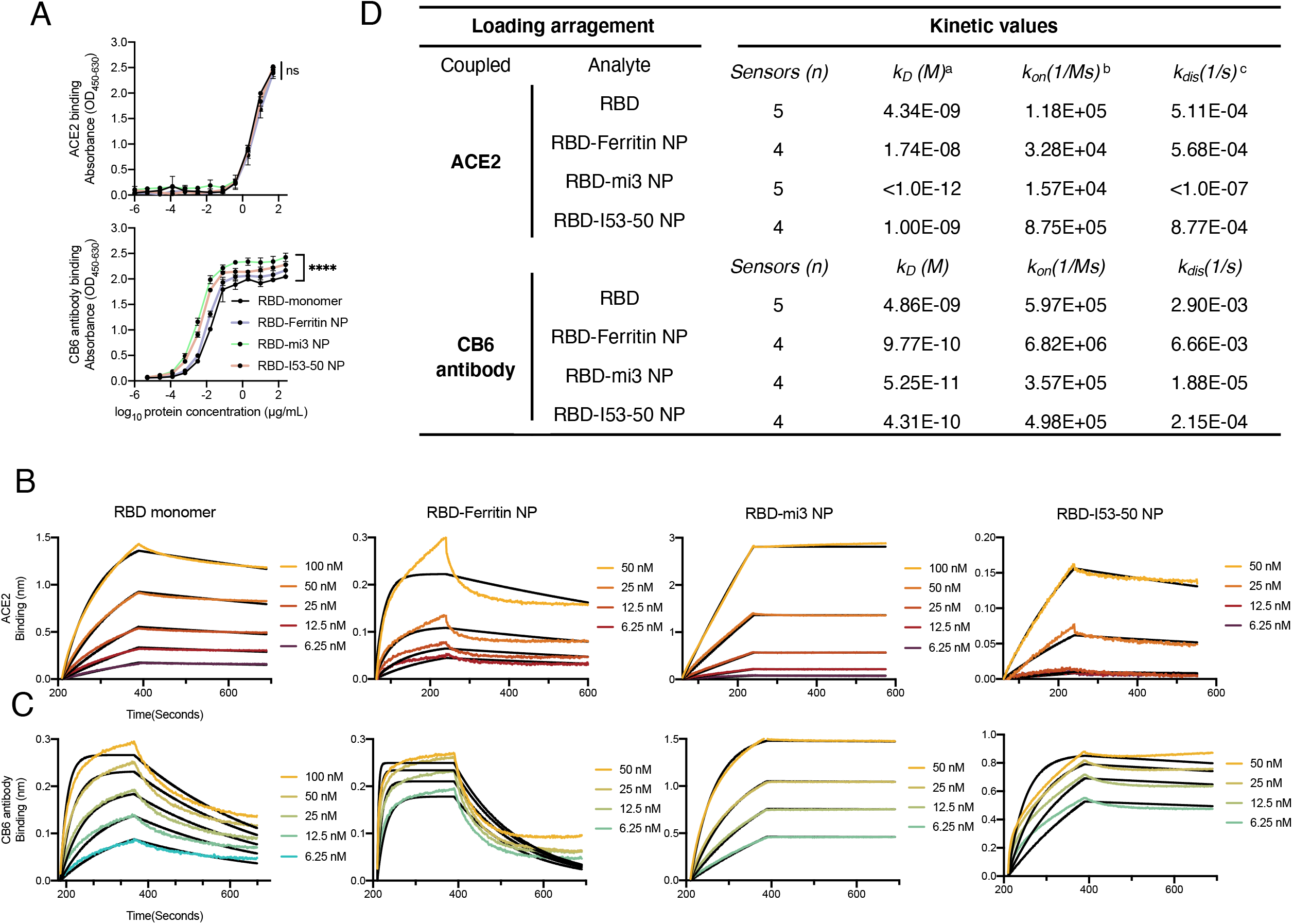
Antigenicity characterization of RBD monomer and RBD-conjugated nanoparticles. (A) ELISA assay of ACE2 and CB6 antibody binding capability. Statistical analysis of binding titers between RBD monomer and the three RBD-NPs was performed using 2-way ANOVA corrected with Dunnett method. (B) (C) Biolayer interferometry (BLI) kinetic assays of RBD monomer and RBD-NPs. (D) Detailed information of BLI assay. a. k_D_: binding affinity constant calculated by k_on_/k_dis_, smaller values generally indicate stronger binding capability b. k_on_: association rates c. k_dis_: dissociation rates

### 2.4. Immunogenicity of RBD-conjugated nanoparticles in BALB/c mice

To compared the immunogenicity of three RBD-conjugated NPs and soluble monomeric RBD, mice were immunized with 5 μg monomeric RBD or corresponding weights of RBD-mi3 NP, RBD-Ferritin NP and RBD-I53-50 NP with equimolar RBD formulated with 50 % (v/v) AddaVax or SAS adjuvant at weeks 0, 2 and 4 (Figure 4A). PBS was used as the negative group. As expected, after prime immunization, almost no binding antibody response was detected in groups immunized with monomeric RBD adjuvanted with both AddaVax and SAS. However, immunized with three RBD-conjugated NPs, RBD-Ferritin, RBD-mi3 and RBD-I53-50, formulated with AddaVax adjuvant elicited 71.8 to 168.4-fold higher binding antibody (ED_50_: 10^3.8±0.4^, 10^3.9±0.2^, 10^4.2±0.2^, respectively) than RBD (ED_50_: 10^2.0^). Analogous to the the Addvax adjuvant, immunized with RBD-Ferritin, RBD-mi3 and RBD-I53-50 adjuvanted with SAS adjuvant induced virtually the same antigen-specific binding antibodies in mice (ED_50_: 10^4.1±0.3^, 10^4.0±0.2^, 10^4.3±0.2^, respectively) after the prime immunization. Next, the RBD-specific antibody titers were substantially increased among groups immunized with monomeric RBD and three RBD-conjugated NPs in the following 1^st^ boost and 2^nd^ boost immunization. ELISA profiles showed that throughout the whole immunization three RBD-conjugated NPs induced significantly higher RBD-specific binding antibody compared to the monomeric RBD, while the titer levels among groups of RBD-conjugated NPs adjuvanted or groups between the two adjuvants were generally similar (Figure 4B). In order to explore the detailed immune response during immunization, we further evaluated the IgG subtype of the antibody elicited and the results showed that a similar trend of antibody titer among different immunization groups could be observed regardless of subtypes of IgG. Moreover, the titer ratio of IgG1:IgG2a among groups were greater than 1 throughout the whole immunization process, indicating a Th2-favored antibody response ^28^ (Figure 4B).

**Figure 4.**
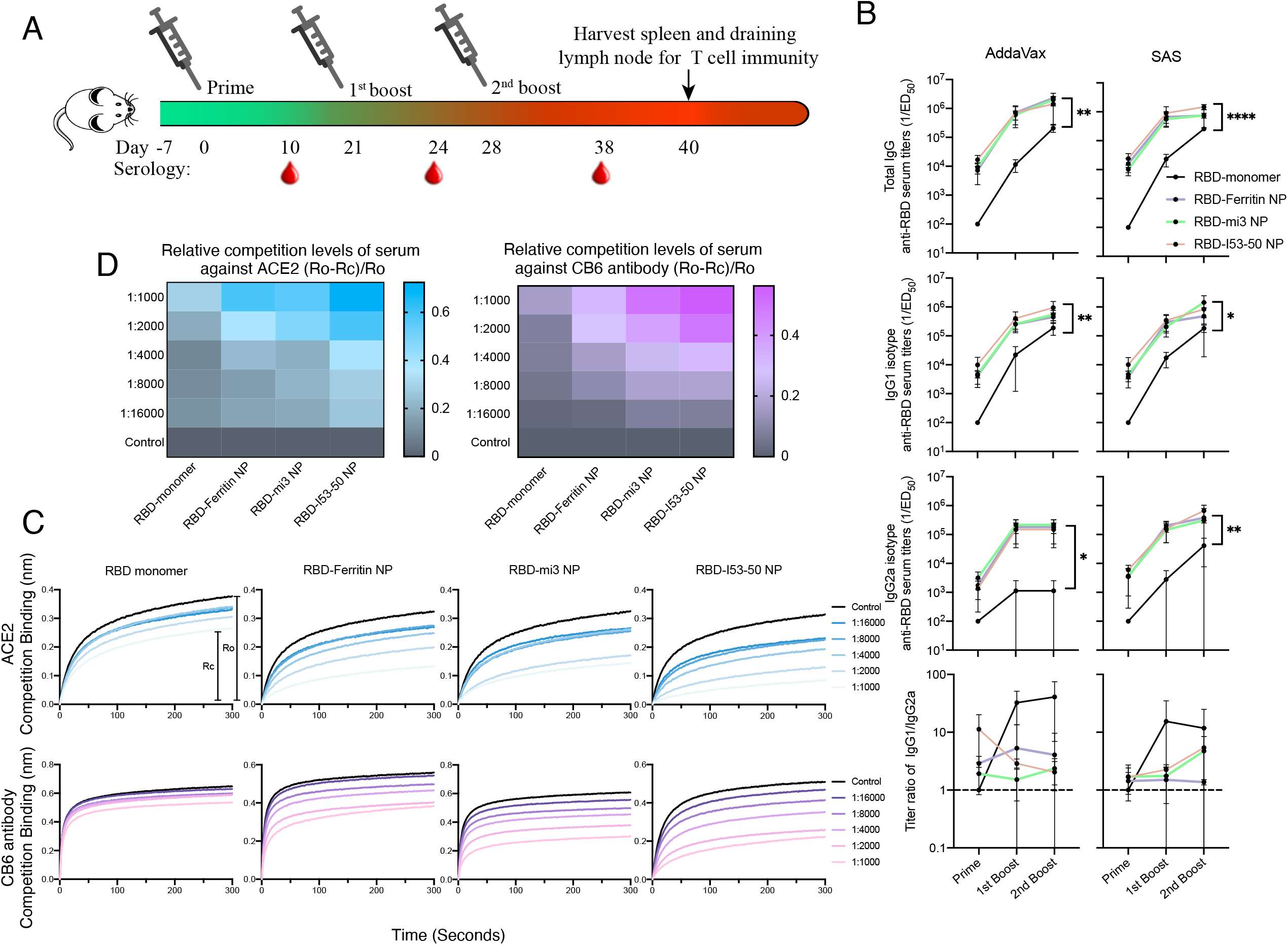
Immunogenicity characterization of RBD monomer and RBD-conjugated nanoparticles. (A) Schematic flow diagram of animal immunization procedures. (B) Serum antibody titers of mice immunized by immunogen adjuvanted with AddaVax or SAS determined by ELISA. Statistical difference between RBD monomer and RBD-NPs are calculated with Two-way ANOVA corrected by Dunnett method with setting the monomer as control group. * p < 0.05; ** p < 0.01; *** p < 0.001; **** p < 0.0001. (C) BLI serum competition assay of sera from immunized with RBD monomer and RBD-NPs adjuvanted with AddaVax against ACE2 or CB6 antibody. Rc represent the binding signal of ACE2 or CB6 under each dilution level. Ro represent the binding signal of serum-free binding signal of ACE2 or CB6. (D) Heatmap overview of competition assay. The competition level presented by ratio (Ro-Rc)/Ro. Brighter color indicates stronger competition against receptor ACE2 or neutralizing antibody CB6 under each dilution level.

Besides the intensity of antibody production, the neutralizing capability of generated antibody would be another critical factor impacting the quality of immunization. Thus, for further evaluation of immunogenicity of RBD-conjugated NPs, serum competition assay through BLI was performed. Sera of mice immunized with antigen adjuvanted with after 2^nd^ boost was retrieved and mixed within each group for a general evaluation. After serial dilution, sequentially-diluted sera were applied to the RBD captured on the biosensors for blocking. It was observed that sera from RBD-conjugated NPs groups can priorly hinder the binding of ACE2 and CB6 antibody to RBD under each dilution level in comparison with the RBD monomer (Figure 4C). As the binding signals were recorded, the non-competing binding curve height Ro and competing binding curve of each dilution level Rc could be used for quantitative analysis. The heatmap of relative competition level of mice sera further displayed that competitive levels of RBD-conjugated NP groups were 4-to 16-fold stronger to the monomeric RBD group (Figure 4D). And as the copies of RBD presented on the surface increased, a stronger competition could be observed when comparing the RBD-Ferritin NP with RBD-I53-50/mi3 NPs. A stronger relative competition level may indicate a more lasting occupation of RBD of the spike protein on the virus intruding and hinder its binding with ACE2 to prevent cell infection, which would further confirmed by neutralizing assays.

Neutralization antibody titers were determined in vitro using SARS-CoV-2 pseudovirus and authentic SARS-CoV-2 virus neutralizing assays. The neutralization titer of sera collected with the RBD-conjugated nanoparticles formulated with AddaVax adjuvant after the 2^nd^ boost immunized mice was ~10-to 120-fold greater than that of sera from the monomeric RBD control group during SARS-CoV-2 pseudovirus-based assay. Analogical result was also observed when immunized with antigen formulated with SAS adjuvant (Figure. 5A). Therefore, we further performed authentic SARS-CoV-2 virus to detected the neutralization activity of sera by method of cytopathic effect (CPE) evaluation and focus reduction test. As shown in Figure. 5B, 90 % focus reduction neutralization antibody titers (FRNT_90_) of post-2^nd^ boost sera from mice immunized with RBD-Ferritin NP, RBD-mi3 NP and RBD-I53-50 NP formulated with AddaVax adjuvant (FRNT_90_:10^4.1±0.5^, 10^4.0±0.6^, 10^4.1±0.3^, respectively) was ~10-to 40-fold higher than monomeric RBD (FRNT_90_: 10^2.1±0.8^). As was similar to AddaVax, three RBD-conjugated nanoparticles adjuvanted with SAS also showed significantly higher FRNT_90_ than RBD monomer (Figure 4B). Otherwise we compared the difference of neutralization activity of sera from all groups after each immunization procedure according the CPE-based microneutralization assay. Results manifested that after the first prime, little neutralizing effect could be observed among all groups. As the immunization procedures progressed, neutralizing effect of sera samples of groups from RBD-conjugated NPs was significantly overwhelming the monomeric RBD group in regardless of adjuvants used. Specially, comparative serum neutralizing activity could be observed after 1^st^ boost for the RBD-conjugated groups while an equal strength of neutralization was postponed to post-2^nd^ boost for monomeric RBD group, during which the neutralizing activities of nanoparticle groups were nearly 10-fold higher. It was interesting to point out that the RBD-Ferritin NP showed a relatively inferior effect compared to the other two nanoparticles when we made a parallel contrast between groups, which was in accordance to the competition assay.

**Figure 5.**
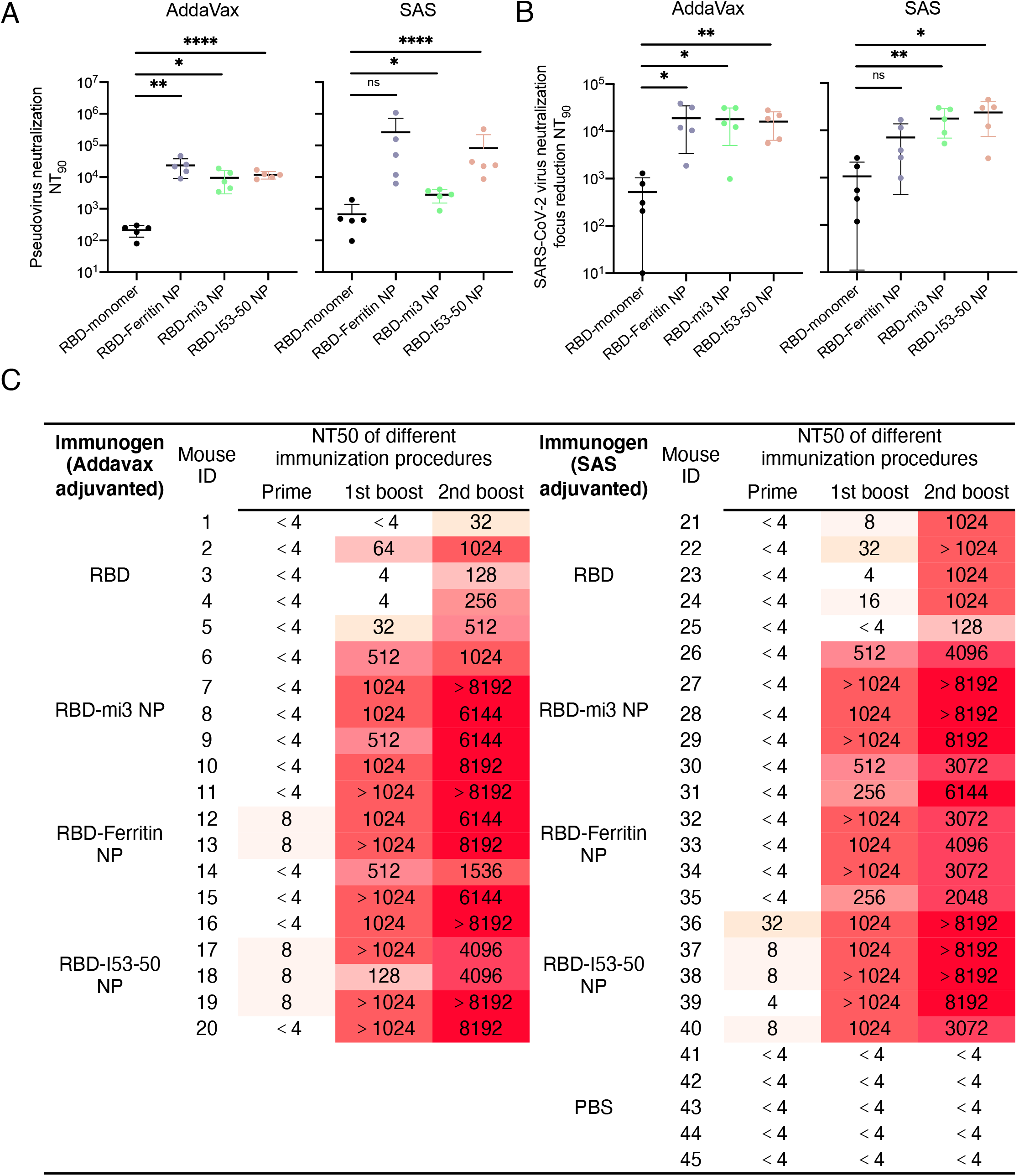
Neutralizing capability of mice sera of RBD monomer and RBD-conjugated nanoparticles. (A) SARS-CoV-2 pseudovirus neutralizing assay shows the NT_90_. (B) SARS-CoV-2 live virus neutralizing assay show the focus reduction NT_90_ (FRNT_90_). Statistical difference of neutralizing titers of mice immunized by immunogen adjuvanted with AddaVax or SAS are calculated with unpaired two-tailed non-parametric Mann-Whitney U test. * p < 0.05; ** p < 0.01; *** p < 0.001; **** p < 0.0001. (C) Table of SARS-CoV-2 live virus neutralizing titers determined by induced cytopathic effect (CPE). Deeper red color represents a higher dilution ratio.

### 2.5. Cellular reponse during the immunization

To explore whether a difference in immune cell level of immune response to antigen manipulate the above effect and determine the T cell immune response elicited by RBD-conjugated NPs, analysis of germinal centers (GCs) B cells, T follicular helper (Tfh) cells and immune cells containing intracellular cytokine was performed. Compared to monomeric RBD, analysis of responding cells in draining lymph nodes following a 2^nd^ boost immunization shown that no significant difference was observed in Tfh and GC cell responses when immunized with three RBD-conjugated NPs (Figure S3A and 3B, Supporting Information). In addition, consistent with the RBD-dimer results as previously reported ^16^, compared with monomeric RBD immunized mice, no substantially increase of T cell responses of collected draining lymph nodes and spleen was detected by flow cytometry in RBD-conjugated NPs immunized mice (Figure S3C and S3D, Figure S4, Supporting Information).

## 3. Discussion

With the worldwide collaboration in SARS-CoV-2 research, various vaccine candidates were raised and validated in preclinical or clinical trials ^29–32^. And to enhance the immunogenicity of antigen used in vaccine, strategies including live-virus platform ^33–34^, viral vector vaccine using vesicular stomatitis virus ^35^ or adenovirus ^36^, lipid nanoparticle-encapsulated mRNA ^37^, or whole inactivated virus ^38–39^ were adopted, among which the multivalent presentation of antigen on nanoparticle protein was regarded as one of the most rapidly-developing method for vaccine design ^40–41^. However, the increased structural redundance during the de novo design of antigen constructed on the multimeric component pulled apart the gap between well-designed blueprint structure and the actual production ^42^. Hence, we reported the design of RBD-based nanoparticles using covalent bond linking strategy as the example for a potent method for rapid antigen-nanoparticle design.

As the global awareness of the urgent need of fast-responding vaccine development grew under the pandemic, a delicate selection of antigen able to elicit competent intensity of neutralizing antibody was demanded. Thus, spike protein as the major viral membrane protein became the focused antigen candidate for vaccine design, leading to selection of different protein subsection used (full ectodomain S protein, S1 segment and RBD) based on the co-structure and functional mate, ACE2 ^8, 13, 43^. Despite a full display of available antigenic site, full length S protein bear uncertainty of prospective immune response due to increasing evidence of versatile mutations ^44^, unpredictable presenting efficacy of neutralizing epitope and antibody-dependent enhancement (ADE) effect ^45^. The shortening of ectodomain of spike protein to S1 maintains a balanced characteristic between full length protein and RBD domain but still carried the inherited shortage from the full-length spike protein ^46^. As the co-structure of RBD and ACE2 has been elucidated, growing attention was put on the RBD as primary antigen for vaccine design and variable strategies were adopted to enhance the immunogenicity including dimerization, nanoparticlization or simple combined use of adjuvant ^12–13^. Among all the strategies, multivalency of antigen and enlargement of antigen size gained most effort due to that increased antigen size and antigen saturation of BCR prolongs the antigen-presentation retention and foster the recognition of antigenic epitope from the immunogen ^47^. To achieve multimeric display of RBD existed as monomer, necessary component was introduced to the original RBD sequence to form commutative bond between RBDs or to add additional scaffold to initiate multimerization, which required skilled structure-guided modification on corresponding antigen and iterations of ideal-to-real test production ^31,48^. The two time-costing prerequisites for fine design of antigen-nanoparticle would expose disadvantages during emergent spread of infectious diseases, especially under pandemic. Therefore, we validated the utilization of covalent bond linking strategy during rapid development of SARS-CoV-2 vaccine and confirmed the viability by comprehensive evaluation.

Ascribing to separation of the expression of antigen RBD and nanoparticles used for antigen capture, the construction and production of proteins could be achieved in different optimal expression systems (Figure 1B). Later the covalent bond linking would be performed easily by incubation of RBD-SpyTag and ΔN1-SpyCatcher-NPs, yielding fully multivalent RBD-conjugated NPs with high structural uniformity and stability, and little sacrifice in assembling efficiency (Figure 1C and 1D, Figure 2). Both monomer RBD and RBD-conjugated NPs underwent further antigenicity inspection and results showed that multivalency RBD-conjugated NPs exhibited stronger affinity to receptor ACE2 and neutralizing antibody CB6 (Figure 3). We immunized Balb/C mice with monomer RBD and RBD-conjugated NPs adjuvanted with AddaVax or SAS. Serum anti-RBD antibody titers of RBD-conjugated NPs were significantly higher than monomer RBD (Figure 4A and 4B) regardless of adjuvant used, indicating the achieved target of nanoparticle design. Neutralizing assay of pseudovirus or authentic SARS-CoV-2 virus proved that elicited neutralizing antibody titers of RBD-conjugated NPs were also far away higher than monomer (Figure 5), which could be explained by the competition assay of immunized mice sera to ACE2 or CB6 antibody (Figure 4C and 4D). Stronger competition behavior from higher dilution level of sera guaranteed a more perfect protection of recognition by RBD from intruding virus. Moreover, it seemed to be creditable that with the increase of valency of RBD upon the nanoparticle surface, a more favorable immunization effect could be induced, as we compared 24-mer RBD-Ferritin NP with the other 2 nanoparticles (Figure 4C and 4D, Figure 5C).

Here, we reported three RBD-based nanoparticle design using a universal strategy in vaccine development and validated three different nanoparticles platform for future need in rapid and general vaccine design. It’s completely viable for replacement of RBD to other antigens from potential risky pathogen as the immunogen core. The independence of antigen screening and expression, nanoparticle scaffold selection, particle assembly and immunogenicity validation would bring helps to researchers devoted to contribute to vaccine development without setting an excess threshold of required structural information and experimental skills, and to manufacturer in commercial production due to a high yield of protein components and shortening process in upper-stream research and development, not only for the current pandemic but for the future battle with unknown pathogens.

## 4. Experimental Section

### Cells and viruses

Vero-E6 (clone E6) and Vero cells were kidney epithelial cells from African green monkey and purchased from ATCC. The HEK293T cell is a human embryonic kidney epithelial cell line and obtained from ATCC. The HEK293F cells were purchased from Life Technologies and maintained in Union 293 medium (Union-Biotech) at 37 °C with 5% CO2 and shaking at 120 rpm. The HEK293T cell expressing human angiotensin-converting nzyme 2, HEK293T-hACE2, is deposited in our lab. All adherent cells were grown in Dulbecco’s modified Eagle’s medium (DMEM) supplemented with 10% fetal bovine serum (FBS) and 1% penicillin-streptomycin at 37 °C with 5% CO2. All cell lines were confirmed free of mycoplasma contamination. In this study, both SARS-CoV-2 strains we used were isolated from COVID-19 patients in Guangzhou (Genbank: MT123290 and GISAID: EPI_ISL_413859).

### Mice

Specific pathogen-free (SPF) six to eight weeks old female BALB/c mice were obtained from the Beijing Vital River Laboratory Animal Technology Co., Ltd. All experimental animal studies were approved by the ethics committee of Sun Yat-sen University Cancer Center (approve number: L102042020000A).

### Gene synthesis and plasmid construction

The SARS-CoV-2 RBD (residue 319-541, GenBank: MN908947) construct for preparation of RBD-based nanoparticle vaccine were human codon-optimized and synthesized by Genscript, and further cloned into mammalian expression vector VRC8400 with a N-terminal Kozak consensus sequence, signal peptide for protein secretion and a C-terminal octa-histidine tag followed by a 13-residues SpyTag ^18^ using the BamHI restriction sites. The SARS-CoV-2 RBD construct for ELISA assay was basically same as the above but without the 13-residue SpyTag. The human ACE2 (residue 19-615, GenBank: NM_021804.2) was synthesized by Genscript and cloned into VRC8400 with a N-terminal Kozak consensus sequence and signal peptide and with a C-terminal octa-histidine tag.

The IgG heavy and light chain variable genes of CB6 mAb (GenBank: MT470196 and MT470197) were human codon-optimized and synthesized by Genscript and cloned into antibody expression vectors.

The ΔN1-SpyCatcher-mi3 construct (GenBank: MH425515) with mutations C76A and C100A based on pentameric I3-01(60-mer) ^17, 24^ was E. coli codon-optimized and synthesized by Genscript and cloned into a modified pET28a* vector with a N-terminal hexahistidine tag. The ΔN1-SpyCatcher was fused to the N-terminal trimeric Ferritin (24-mer) ^23^ or trimeric I53-50A1.1PT1 (60-mer) ^25^ using the (GGS)4 spacer, and E. coli codon-optimized and synthesized and then cloned into a modified pET28a* vector with a N-terminal hexahistidine tag or a C-terminal hexahistidine tag, respectively, by Genscript.

The I53-50B.4PT1 was codon-optimized and synthesized and then cloned into a modified pET28a* vector with a C-terminal hexahistidine tag by Genscript.

### Protein expression and purification in HEK293F cells

The expression plasmid was transformed into DH5a competent cell for plasmid DNA extraction using the NucleoBond Xtra Maxi kit according to the manufacture protocol. The SARS-CoV-2 RBD monomer and hACE2 proteins were produced in HEK293F cells. The HEK293F cells were cultured in Union 293 medium at 37 °C, 80-90% humidity, 5% CO2 with rotation at 120 rpm for expansion. Then cells were transiently transfected with 2mg expression plasmid per 1 liter using the Polyethylenimine (PEI) MAX (Polysciences) at a density of 1.0 x 106 cells/ml in fresh Union 293 medium. After 5 days culture, the cell culture was collected and centrifuged at 4°C, 8000 g for 1h. Collected supernatant was further filtered using Steritop (0.22 μm pore size; Guangzhou Jet Bio-Filtration Co., Ltd) and concentrated to 1/10 volume using tangential flow filtration system (10 kD retention molecular weight, Millipore). Then concentrated supernatant was purified by immobilized metal-affinity chromatography with Ni-NTA resin (Roche) stocked in WET FRED gravity flow columns (IBA) and beads were eluted with buffer composed of 50 mM HEPES, pH 7.3, 300mM imidazole and 300 mM NaCl. The eluate was concentrated and further purified with size exclusion chromatography using Superose 6 Increase 10/300 GL gel filtration column (GE Healthcare) in a buffer composed of 50 mM HEPES, pH 7.3 and 300 mM NaCl. Fractions of the target peak were and pooled and concentrated using centrifuge tubes (10KDa MWCO, Millipore), and followed by store in 4 °C for further use.

CB6 antibody was expressed and purified as previously reported ^27^. Briefly, the plasmids encoding the IgG heavy and light chain gene were transiently tansfected into HEK293F cells at a ratio of 5:6. The supernatant of culture was collected 5 days after transfection, centrifuged and purified by affinity chromatography with protein A resin (Genscript), and subjected to desalt in a composed buffer 50 mM NaPO4, 150 mM NaCl, pH 7.3 using HiTrap Desalting column (GE Healthcare) with AKTA pure chromatography system (GE Healthcare).

### Protein expression and purification in E. coli

The modified pET28a* expression plasmid of Δ N1-SpyCatcher-mi3, ΔN1-SpyCatcher-Ferritin, Δ N1-SpyCatcher-I53-50A1.1PT1 and I53-50B.4PT1 was transformed into Rosetta™ (DE3) competent cells (TIANGEN). After incubation for overnight at 37 °C on TB-agars culture plate supplemented with 50 mg/mL kanamycin and 33 mg/mL chloramphenicol, a single positive colony was selected and inoculated into 10 mL TB medium in the presence of 50 mg/mL kanamycin and 33 mg/mL chloramphenicol and was grown overnight at 37 °C with shaking at 220 rpm. The culture was added to the 3L baffled triangle shake flasks containing 1 L TB medium and 50 mg/mL kanamycin, and grown at 37 °C with shaking at 150 rpm. When OD600 value of the culture reached up to 0.6~0.8, isopropylthiogalactoside (IPTG) was added to a final concentration of 1 mM for induction at 20 °C for 16-20 h with shaking at 150 rpm. The bacterial cultures were harvested and centrifuged at 20 °C, 2450 g for 15 min

For ΔN1-SpyCatcher-mi3 purification, cell pellets were resuspended in lysis buffer (250 mM Tris, pH 8.5, 300 mM NaCl, 30 mM imidazole, 1 μM DNases, 0.75% CHAPS, 5 mM MgCl_2_ and EDTA-free protease inhibitor cocktail [Roche]), and lysed with high pressure cell homogenizer (Union-Biotech) at a pressure of 800 MPa. The suspensions were centrifuged for supernatant collection, filtered with Steritop (0.22 μm pore size), and purified with the gravity flow columns containing Ni-NTA resin. Beads were eluted with 50 mM HEPES, pH 8.0, 300 mM NaCl, 300 mM imidazole and 0.75% CHAPS, and the elution was concentrated to 1mL and loaded onto the size exclusion chromatography using Superose 6 Increase 10/300 GL gel filtration column (GE Healthcare) pre-equilibrated with 50 mM HEPES, pH 8.0 and 300 mM NaCl. Peak fractions were identified with SDS-polyacrylamide gel electrophoresis (PAGE) analysis to determine whether target protein was collected. After a concrete confirmation of purity and yield of protein, the fractions were pooled, concentrated, and stored at 4 °C.

For Δ N1-SpyCatcher-Ferritin and Δ N1-SpyCatcher-I53-50A1.1PT1 proteins purification, cell pellets were resuspended in lysis buffer (50 mM HEPES, pH 7.3, 300 mM NaCl, 30 mM imidazole, 1mM DTT, 0.75% CHAPS, 1 μM DNases, 5◻mM MgCl_2_ and EDTA-free protease inhibitor cocktail [Roche]). Purification was similar as the above instead of the elution buffer used (50 mM HEPES, pH 7.3, 300 mM NaCl, 300 mM imidazole, 1mM DTT and 0.75% CHAPS), and equilibration buffer for SEC (50 mM HEPES, pH 7.3 and 300 mM NaCl).

Endotoxin of all purified proteins was removed with ToxinEraserTM Endotoxin Removal Kit (Genscript) in accordance to the manufacturer’s instruction. The remnant endotoxin was identified with ToxinSensor^TM^ Chromogenic LAL Endotoxin Assay Kit (Genscript) and no more than 0.1 EU/mL of endotoxin was detected.

### Preparation of RBD-conjugated nanoparticles

To assemble the RBD-conjugated nanoparticles, RBD-SpyTag should be incubated with ΔN1-SpyCatcher-NPs to form a covalent peptide bond in between due to an automatic reaction of SpyTag-SpyCater system (Banerjee and Howarth, 2018; Bruun et al., 2018). Purified Δ N1-SpyCatcher-Ferritin or ΔN1-SpyCatcher-I53-50A1.1PT1 protein presented at a subunit concentration of 50 μM were incubated in a 1:8 molar excess ratio with RBD-SpyTag for overnight at room temperature buffered with 50 mM HEPES, pH 7.3, 300 mM NaCl. For ΔN1-SpyCatcher-mi3, 50 μM subunit of concentration of protein was mixed in a 1:8 molar excess ratio with RBD-SpyTag for overnight at room temperature buffered with 50 mM HEPES, pH 7.5, 300 mM NaCl, 5% glycerol. In order to separate the conjugated nanoparticles with the empty-NPs and monomer RBD, all incubated substrates were applied to size exclusion chromatography using Superose 6 Increase 10/300 GL gel filtration column pre-equilibrated with 50 mM HEPES, pH 7.3, 300 mM NaCl. Fraction were collected for SDS-PAGE analysis. The protein of interest was selected, concentrated and stored at 4 °C.

### SDS-PAGE analysis

SDS-PAGE was performed as previously described ^49^. Briefly, five micrograms of purified protein by added the 5x loading buffer were heated at 95 °C for 5 min, and loaded on 12 % Tris-glycine gels for 30 min at 300 V. Gels were stained with coomassie brilliant blue (Beyotime, China) and destained with 30% methanol,10% glacial acetic acid in double distilled water and subjected to film by ChemiDoc systerm (BioRad).

### Dynamic light scattering

DLS was carried out to characterize the hydrodynamic diameter and polydispersity index of RBD-conjugated nanoparticles and individual nanoparticles using Zetasizer Ultra instrument (Malvern Panalytical). Briefly, purified proteins were centrifuged at 16,250 g, 4 °C for 10 min to remove any aggregates, and diluted to at a concentration of 0.5 mg/mL in PBS and loaded onto the disposable solvent resistant micro cuvette.

The particle distribution of the purified protein was determined by measuring the intensity of the light scattered by the sample using the avalanche photodiode detector placed at a measurement angle of 173° at 25 °C. Each sample was measured in triplicate and the average values of the hydrodynamic diameter and polydispersity index of the sample were recorded and analyzed using the manufacturer’s software (Malvern Panalytical).

### Negative stain electron microscopy

Approximately 5 μL aliquot of purified nanoparticle at a concentration of 0.05-2 mg/mL was applied to freshly glow-discharged 300-mesh copper grids and incubated for 1 min. Excess liquid were blotted with filter paper. The grids were washed twice by double distilled water and blotted, and then negatively stained freshly 2 % (w/v) uranyl acetate for 45s, and followed by air dried. Grids were imaged with FEI Tecnai T12 transmission electron microscope (FEI, USA) operating at 120 kV. The digital micrographs were obtained at 150,000x magnification.

### Nano differential scanning fluorimetry

NanoDSF Systems were conducted to measure the thermostability and aggregation of purified protein using Prometheus NT.48 instrument (NanoTemper Technologies). A 10 μL aliquot of sample diluted to a concentration of 5 mg/mL were applied to quartz capillary cassette and placed into card slot. The scan temperature was increased linearly starting from 20.0 □ to 95 □ at a scan rate of 1 °C. The thermal transition midpoint (T_m_) and aggregation results from start to finish (Tagg) were reported and analyzed in PR.ThermControl software (NanoTemper Technologies). Three replicates are measured for each sample.

### Enzyme-linked immunosorbent assay

ELISA assay was performed to examine the binding ability of purified protein to the receptor ACE2 and SARS-CoV-2 RBD-specific CB6 Antibody. Purified RBD monomer and RBD-conjugated NPs was diluted to at a concentration of 1 μg/mL and precoated on 96-well microplates (Corning) (100 μL/well) in triplicate overnight at 4 □. The plates were washed three times with PBS with 0.05 % Tween 20 (PBS-T), and blocked with ELISA blocking buffer (2 % gelatin, 5 % casino and 0.1 % proclin 30 in PBS) for 1 h at 37 □. The plates were then incubated with sequentially 1:5 diluted ACE2 starting from 5×10^-4^ ng/mL to 10^-2^ ng/mL in ELISA blocking buffer. After 1 h incubation, the plates were washed with PBS-T for five times, and 100μL hACE2-specific rabbit antibody (Sino Biological Inc) diluted at a ratio of 1:5000 was added to wells at 37 □ for 1 h. The plates were washed with PBS-T 5 times and a 1:5000 dilution of HRP-conjugated goat anti-rabbit IgG antibody (Promega) was added for 45 min at 37 □. After the final round of wash to remove all disassociated antibody, substrate 3,3′,5,5′-Tetramethylbenzidine (TMB, Sangon Biotech) was added to generate chromogenic reaction for 15 min at room temperature, and the reaction would be suspended by a followed addition of 2M H_2_SO_4_. The well-absorbance at 450 nm and 630 nm was immediately recorded by SpectraMax Plus plate reader (Molecular Devices, USA).

As for the assay of CB6 antibody binding, the coating, incubating and chromogenic reaction generating were similar to the ACE2 assay instead of that primary antibody binding to antigen was not required and that HRP-conjugated goat anti-human IgG antibody (Promega) would be used as the secondary antibody to detect binding of CB6 antibody to the antigens.

### Bio-layer Interferometry

BLI analysis were performed on an Octet Red 96 (Fortebio) instrument at 30 °C with shaking at 1000 rpm. Signals were collected at a standard frequency as default (5.0 Hz).

#### Kinetic assay

Firstly, the Streptavidin (SA) biosensors (Fortebio) were pre-incubated in PBS (ThermoFisher) containing 0.05 % Tween 20 (Sigma-Aldrich), the assay buffer used throughout the whole procedures, for 15 minutes. To couple the RBD protein on the biosensors, EZ-link-Sulfo-NHS-biotin biotinylation kit (ThermoFisher) was used to biotinylate the RBD/ACE2/CB6 antibody in the following of the instruction as below.

Step1: Calculation of the amount of biotinylation agent used

1. Calculate millimoles of biotin reagent to add to the reaction for a 20-fold molar excess:

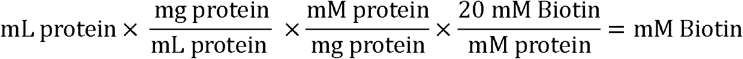

20 = Molar fold excess of biotin
2. Calculate microliters of 10 mM biotin reagent solution to add to the reaction:

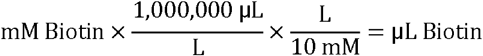

Step2: Biotinylating

Add the calculated amount of 10 mM biotin reagent into the RBD/ACE2/CB6 protein in PBS (5 mg/mL, 200 μL) and incubated the reaction system at room temperature for 30 minutes.

Step3: Desalting

Equilibrate the desalting column PD-10 (GE Pharmacia) with 10 mL of PBS. After the equilibration, the reaction solution would be added in the column and then washed and eluted with 400 μL PBS separately.

To perform kinetic assay, after 60 s baseline, ACE/CB6-biotin protein diluted with the buffer was captured on the SA sensor at 2 μg/mL for 120 s. Then 2 fold-diluting RBD-copy molar concentrations of RBD monomer or different RBD-conjugated NPs were associated to the biosensor for 180 s, followed by a 300 s disassociation and 3 rounds of regeneration with 10 mM Glycine pH 1.5. The curve data was analyzed by ForteBio data analysis software. Raw curves aligned at association were adjusted with baseline signals before a 1:1 binding model fitting was performed. Then a global fit to all binding curves was conducted to render overall kinetic parameters (kD, kon, kdis etc.)

#### Serum competition assay

Mice serum from AddaVax-adjuvanted immunization group harvested after 2^nd^ boost were collected and equal volume (5 μL) of sera from each mouse within the same immunogen group was mixed together in representative of overall characteristic of the group. To perform the competition assay, RBD-biotin protein was captured on the biosensor as the above at 5 μg/mL. Then to saturate the RBD, 2 fold-diluting mice sera mixtures from each group and control PBST were loaded to the biosensor for 300 s. After the end of sera loading, 400 mM ACE2 or CB6 antibody were associated to the biosensors for 300 s to detect the competitive binding signal under the saturation of each dilution level of mice sera. Sensors were also regenerated with 10 mM Glycine pH 1.5. Real-time signal data was collected and the competition behavior was displayed by the ACE2/CB6 binding signal of different curves. Binding signal data were retrieved from curve, Ro represented the saturated non-competing binding curve height and Rc represented saturated competing binding curves of each dilution level. Relative competition levels of each serum dilution level could be calculated as (Ro-Rc)/Rc.

### SARS-CoV-2 pseudovirus production

The SARS-CoV-2 pseudovirus were produced as previously described ^50^, with some modification. Briefly, gene encoding SARS-CoV-2 S (GenBank: QHU36824.1) with a 19-amino-acids deletion in C-terminal was human codon-optimized and cloned into the expression vector pCMV14-3×Flag, a generously gift from Zhaohui Qian, Chinese Academy of Medical Sciences. HEK293T cells were co-transfected with the plasmids of PsPAX2, pCMV14-SARS-CoV-2 S ∆CT-3×Flag and pLenti-GFP at a ratio 1:2:1 using PEI-MAX (mentioned above). After 5 hours, the supernatant was replaced with fresh DMEM supplemented with 10% FBS. Sixty hours post-transfection, the supernatant containing SARS-CoV-2 pseudovirus were harvested and centrifuged at 4 °C, 3000 g for 10 min to remove cell debris, followed by a filter process with Steritop (0.45 μm pore size, Guangzhou Jet Bio-Filtration Co., Ltd). Sterile PEG-8000 solution was added to the clarified supernatants and the solution was incubated for 1 hour at 4 °C. The mixture was then centrifuged, concentrated, and resuspended with DMEM for final collection of viruses. The storage of virus was at −80 °C.

### BALB/c mice immunization

Forty-five 6-8 weeks-old female BALB/c mice were purchased from the Beijing Vital River Laboratory Animal Technology Co., Ltd, and arbitrarily divided into 9 groups. Before immunization, purified immunogen was diluted with PBS and gently formulated with an equal volume of AddaVax™ adjuvant (InvivoGen) or Sigma Adjuvant System (SAS, Sigma), and incubated overnight to achieve full absorption of antigen upon the surface of adjuvant particles at 4 °C with shaking at 40 g. Each group of mice received three immunizations at weeks 0, 2 and 4 via a subcutaneous route. The immunization dose was 5 μg of RBD monomer, or corresponding weights of RBD-conjugated nanoparticle immunogens containing equal molar of RBD as the monomers which were RBD-mi3 (9.51 μg), RBD-Ferritin (9.34 μg) and RBD-I53-50 (11.91 μg). PBS were served as a negative control. Blood samples were harvested 10 days following each immunization, and were placed at 37 °C for 30 min to reach ample coagulation. Then blood samples were centrifuged at 16,250 g, 4 °C for 10 min and the upper layer serum was gently extracted, heat-inactivated at 56 °C for 30 min to deactivate the complement factors and pathogens, and then stored at −20 °C for future analysis.

### Serum ELISA

1 μg/mL RBD monomer without SpyTag in PBS was precoated on 96-well maxiSorp ELISA microplates (Corning) overnight at 4 [. Plates were blocked with ELISA blocking buffer for 1 h at 37 □. Mouse serum samples were 5-fold sequentially diluted with blocking buffer starting at 1:50, and were added to the coated plates, and incubated at 37 °C for 1 h. After incubation, the plates were washed with PBS-T five times and added with a 1:5000 dilution of HRP-conjugated goat anti-mouse IgG antibody (Promega) in blocking buffer at 37 °C for 45 min. For analysis of RBD-specific IgG isotype titers, a 1:5000 dilution of HRP-conjugated goat anti-mouse IgG1 and IgG2a antibody was incubated at this step. Plates were washed, colored by TMB, and quenched with H_2_SO_4_. The absorbance at 450 nm and 630 nm of each well was immediately determined by SpectraMax Plus plate reader (Molecular Devices, USA). Results were plotted and fitted by GraphPad Prism 8 and EC_50_ value were calculated using 4-parameters nonlinear regression fitting from fitted curve.

### Pseudovirus-based neutralization assay

Pseudovirus-based neutralization assay was performed to evaluate the neutralization ability of the sera by immunized mice. In brief, approximately 1.75×10^4^ HEK293T-hACE2 cells were seeded onto 96-well culture plates overnight at 37 °C in 5 % CO2. Sera were 4-fold serially diluted with complete DMEM medium starting at 1:20, and pre-incubated with an equal volume of pseudovirus of SARS-CoV-2 at 37 °C for 2 h. The mixtures were then added to the HEK293T-hACE2 cells for infection. After 2 hours incubation, the mixtures were replaced with fresh DMEM medium containing 2 % FBS and 1 % penicillin and streptomycin, and incubated for 48 hours at 37 °C, 5 % CO2. Wells treated with only medium or virus without incubation with serum were set as negative control and positive control in each plate, respectively. Afterwards, the cells were lysed with lysis buffer and luciferase activity was immediately measured by dual-glo luciferase assay system (Promega). The inhibition rate was calculated as (sera signals – blank control signals)/ (virus signals – blank control signals) * 100 %. The 90 % neutralization antibody titers (NT_90_) were determined using 4-parameter s nonlinear regression fitting from fitted curve using GraphPad Prism 8.

### Authentic SARS-CoV-2 virus-based neutralization assay

All of microneutralization assay for authentic SARS-CoV-2 virus we used in this study were performed in a BSL-3 facility. Two methods, the authentic SARS-CoV-2 virus-induced cytopathic effect (CPE) and focus reduction neutralization test (FRNT) were used to evaluate the neutralizing antibody titers of sera from immunized mice. Briefly, sera were 4-fold serially diluted starting at 1:4 with DMEM supplemented with 2 % FBS and 1 % penicillin and streptomycin, and mixed with the equal volumes of 100 half tissue culture infective doses (100 TCID_50_) SARS-CoV-2-virus of 2020XN4276 strain at 37 °C for 2 h. Afterwards, the sera-virus mixture was added to pre-plated Vero-E6 cells in 96-well culture plate, and incubated for an additional 96 h at 37 °C in 5 % CO_2_ to observe the CPE at 40X magnification. Wells with pure virus treated, pure diluted sera treated or cell only was set as controls for each plate. Virus back titration was performed in each plate. All diluted serum samples were tested in duplicate. The neutralization antibody titers of all of sera were defined as the reciprocal of serum dilution that could neutralize 50 % of virus infection at 4 days post-infection.

As for the FRNT method, serum samples were 5-fold serially diluted starting at 1:10, and mixed with the equal volumes of 100 focus forming unit (FFU) SARS-CoV-2 virus of human CHN/IQTC01/2020 strain in 96-well culture plate, and incubated for 1 h at 37 °C. The mixtures were then added to the 96-well plates that pre-seeded with Vero-E6 cells. After incubation for 1 h at 37 °C, 5 % CO2, mixtures were removed and replaced with 100 μL MEM containing 1.2 % carboxymethylcellulose pre-warmed to 37 °C for an additional 24 hours culture. Thereafter, cells were fixed with 4% paraformaldehyde and permeabilized with 0.2% Triton X-100 in PBS, and then incubated with rabbit anti-SARS-CoV-2 nucleocapsid protein antibody (Sino Biological, Inc) for 1 hour at ambient temperature, followed by adding of a 1:4000 dilution of HRP-conjugated goat anti-rabbit IgG antibody (Jackson ImmunoResearch Laboratories, Inc. West Grove, PA). Plates were colorated using TrueBlue™ Peroxidase Substrate (KPL). Foci were counted on an ELISPOT reader (Cellular Technology Ltd. Cleveland, OH). The 90 % neutralization antibody titers (NT_90_) were defined as the reciprocal of serum dilution that could inhibit 90 % FFU of virus-infection, and was calculated with a 4-parameter nonlinear regression fitting from fitted curve using GraphPad Prism 8.

### Germinal center and Tfh analysis of mice drained lymph nodes

Immunized mice were sacrificed by CO_2_ inhalation 12 days after the 2^nd^ boost immunization (40 days).To simultaneously identify germinal center B cells and T follicular helper cells of mice drained lymph nodes, cell suspensions of draining lymph nodes from mice sacrificed were stained with fixable viability stain 780 (BD Biosciences) and blocked with anti-CD16/32 antibody (BD Biosciences), followed by labeling with anti-B220-BV421 (BD Biosciences), anti-IgD-PE (BD Biosciences), anti-GL7-Alexa Fluor 647 (BD Biosciences), anti-CD95-FITC (BD Biosciences), anti-CD4-BV510 (BD Biosciences), anti-CD44-BV786 (BD Biosciences), anti-ICOS-PE-Cyanine7 (BD Biosciences), anti-CXCR5-PE-CF594 (BD Biosciences) and anti-PD-1-APC-R700 (BD Biosciences) in PBS in the presence of 2 % BSA. The fluorescence signal of labeled samples was acquired on a CytoFLEX S flow cytometry (BECKMAN COULTER).

### Intracellular cytokine staining

Drained lymph nodes and spleen were harvested and washed with RPMI 1640 medium. Then tissues were abraded into cell suspensions by the piston handle of 2 ml syringe in the culture medium (RPMI 1640 containing 10 % FBS and 1 % antibiotics). The suspension was filtered with 40 μm nylon mesh cell strainer (Sangon Biotech). Cells were washed with culture medium and sterile erythrocyte lysis buffer was added (1.5 M NH_4_Cl, 100 mM NaHCO_3_, 10 mM EDTA in deionized water, pH 7.4) to remove red blood cells, followed by staining with fixable viability stain 780 (BD Biosciences) for 30 min at ambient temperature. Approximately 1.0 x10^6^ cells were added to the 6-wells plates and treated with anti-CD16/32 antibody (BD Biosciences) to block the Fc receptor, and then stimulated with 15 μg/mL purified RBD monomer for 3 h at 37 °C. After incubation, GolgiStop and GolgiPlug (BD Biosciences) was added to each well for an additional 15 hours at 37 °C. Next, the cells were harvested and washed twice with culture medium, and labeled with anti-CD3e-PerCP-Cy5.5 (BD Biosciences), anti-CD4-BV510 (BD Biosciences) and anti-CD8a FITC (BD Biosciences) in PBS in the presence of 2 % BSA, after which cells were further fixed with 4 % paraformaldehyde and permeabilized with permeabilization buffer (2 % BSA, 0.1 % saponin, 0.05 % Na_3_N in PBS). Finally, cells were washed with PBS in the presence of 2 % BSA and incubated with anti-IFN-γ-PE-CY7 (BD Biosciences), anti-IL-2-APC (BD Biosciences), anti-TNF-α-PE (BD Biosciences) and control anti-IgG1 antibody for 30 min at 4 °C. The fluorescence signal of labeled samples was acquired on a CytoFLEX S flow cytometry (BECKMAN COULTER).

### BALB/c mice challenge

For SARS-CoV-2 challenge experiment, the female 6-8 weeks old mice were arbitrarily divided into 5 groups in each group. All purified antigens were prepared by mixed 100 μL protein solution diluted in PBS with an equivoluminal AddaVax adjuvant. Groups of twenty mice were immunized subcutaneously with a total protein dose corresponding to 10 μg of the RBD antigen on week 0 and 3. Purified gp350D123 protein of Epstein-Barr virus, including the RBD, formulated with AddaVax adjuvant was used as negative control. Blood were collected at week 2 and 5 for analysis. Sixty days after the second immunization, the mice were lightly anesthetized with isoflurane and intranasally transduced with 2.5x 108PFU of Ad5-hACE2 virus. Five days following transduction, the transduced mice were challenged with 1×105 PFU of SARS-CoV-2 via the intranasal route. Weight changes of the challenged mice were observed for ten consecutive days. At 1 and 3 days after challenge, 4 mice in each group were sacrificed and their lung tissues were collected for titration of the virus titers. On day 4 after challenge, 2 mice in each group were sacrificed and necropsied, and lung tissues were collected for histopathological analysis.

### Sequence alignment and analysis

Except the sequence of Wuhan-Hu-1 which was first identified from a COVID-19 patient in Wuhan city ^5^, obtained from National Center for Biotechnology Information (NCBI) database, other sequences are obtained from the Global Initiative on Sharing All Influenza Data (GISAID). The accession numbers of the RBD sequences of representative SARS CoV-2 strains isolated in different countries are as follows: Wuhan-Hu-1 (Genbank: MN908947), South_Korea/KCDC2489/2020 (EPI_ISL_514892), Thailand/NIH-2492/2020 (EPI_ISL_430841), Japan/Hu_DP_Kng_19-027/2020 (EPI_ISL_412969), India/OR-RMRC25/2020 (EPI_ISL_455308), USA/WA-UW-1762/2020 (EPI_ISL_424245), Mexico/CMX-IMSS_01/2020 (EPI_ISL_424731), Canada/ON_PHL2294/2020 (EPI_ISL_418384), Australia/VIC546/2020 (EPI_ISL_426809), Greece/218_35009/2020 (EPI_ISL_437886), Greece/218_35009/2020 (EPI_ISL_437886), Greece/218_35009/2020 (EPI_ISL_437886), Greece/218_35009/2020 (EPI_ISL_437886), Russia/SCPM-O-08/2020 (EPI_ISL_451970), Spain/Madrid_LP24_5999/2020 (EPI_ISL_428680), Sweden/20-50261/2020 (EPI_ISL_469078), Switzerland/ZH-1000477102/2020 (EPI_ISL_413019), Portugal/PT0533/2020 (EPI_ISL_454257), Scotland/CVR138/2020 (EPI_ISL_425681), Denmark/SSI-101/2020 (EPI_ISL_415646), England/20134020004/2020 (EPI_ISL_423108) and Iceland/348/2020 (EPI_ISL_424372), Nigeria/OS085-CV14/2020 (EPI_ISL_455424) and Venezuela/VEN-95072/2020 (EPI_ISL_476704).

### Quantification and statistical analysis

Kinetic parameters of Biolayer interferometry was rendered by Octet Data Analysis software (Fortebio), and detailed curve fitting method could be found in methods. Statistical analyses of all experimental results were performed with GraphPad Prism 8.01 software. Except the results of flow cytometry are expressed as a percentage of positive cells, all of results are presented as mean ± SEM. Method used for statistical difference between groups could be found in figure legends or corresponding methods for details.

## Acknowledgements

This study was supported by the Sun Yat-sen University “Three major” scientific research special projects in 2020 (No. 84000-31143412), the National Natural Science Foundation of China (No. 81801645, 81830090, 81520108022, 81702001), the China Postdoctoral Science Foundation (No. 2017M612818), the National Science and Technology Major Project (No. 2018ZX09739002-004), the National Key Research and Development Program (2017YFA0505600, 2016YFA0502101), the Natural Science Foundation of Guangdong Province (No. 2017A030312003), the Guangdong Province Key Research and Development program (No. 2019B020226002), the Guangzhou Science Technology and Innovation Commission (No. 201607020038),.

## Author contributions

Y.K., C.S., Z.Z., R.Y., C.K., J.Z.,and M.Z. conceived and designed the project; M.Z. supervised the project; Y.K., C.S. and M.Z. wrote and edited the manuscript; Y.K. purified the RBD-conjugated NPs protein and performed the DLS and DSC experiment; Y.K. and Q.Z. performed the negative-stain EM; Y.K. and C.S. performed the mice experiment and analyzed the results; X.C. and C.S. performed the BLI assay; Y.K., R.Y., P.Z. and Z.Z. performed the neutralization assay.

## Competing interests

Mu-Sheng Zeng has filed patent applications for the development of SARS-CoV-2 RBD-conjugated nanoparticle vaccine candidate. The authors declare no competing financial interests.

---

**Figure S1.**
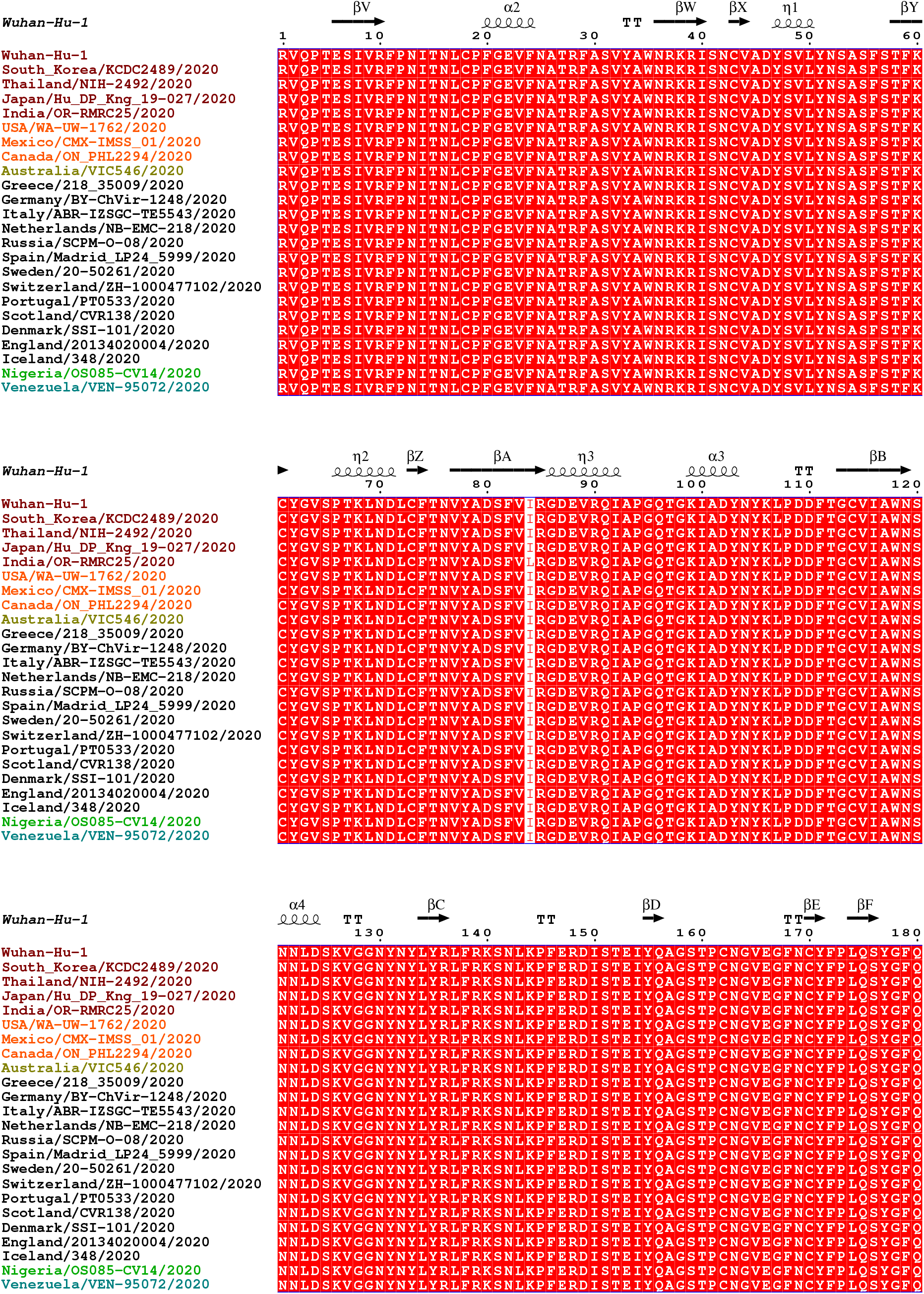
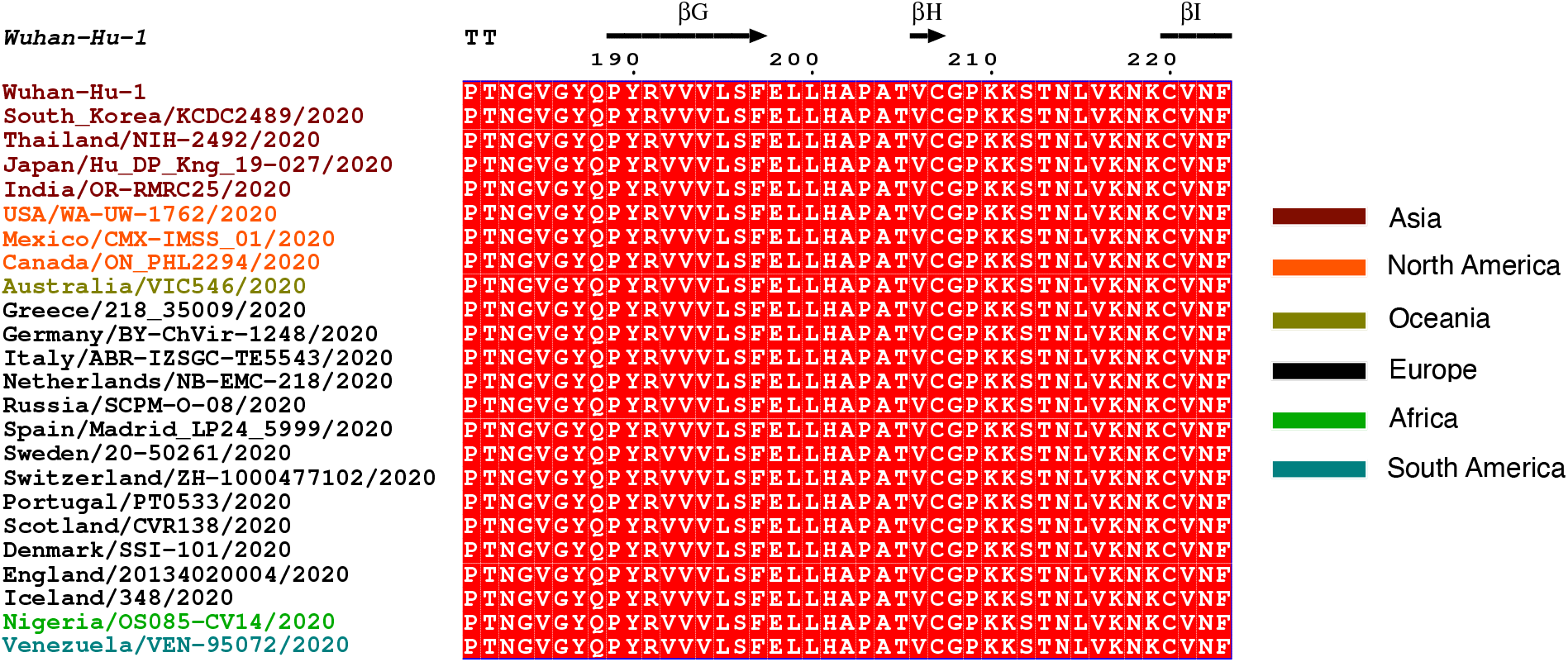
Sequence alignment of RBD from 24 representative SARS-CoV-2 strains isolated from six continents. The sequence of 24 RBD was downloaded from Genbank and GISAID. The strain isolated from Asia, North America, Oceania, Europe, Africa and South America was colored by brown, orange, olivedrab, black, green and cyan, respectively. Conserved residues are highlighted in red. Multiple sequences were aligned by MAFFT (https://mafft.cbrc.jp). The sequence alignment was converted with Clustal X2 (http://www.clustal.org) and visualized with ESPript 3.0 ^51^.

**Figure S2a.**
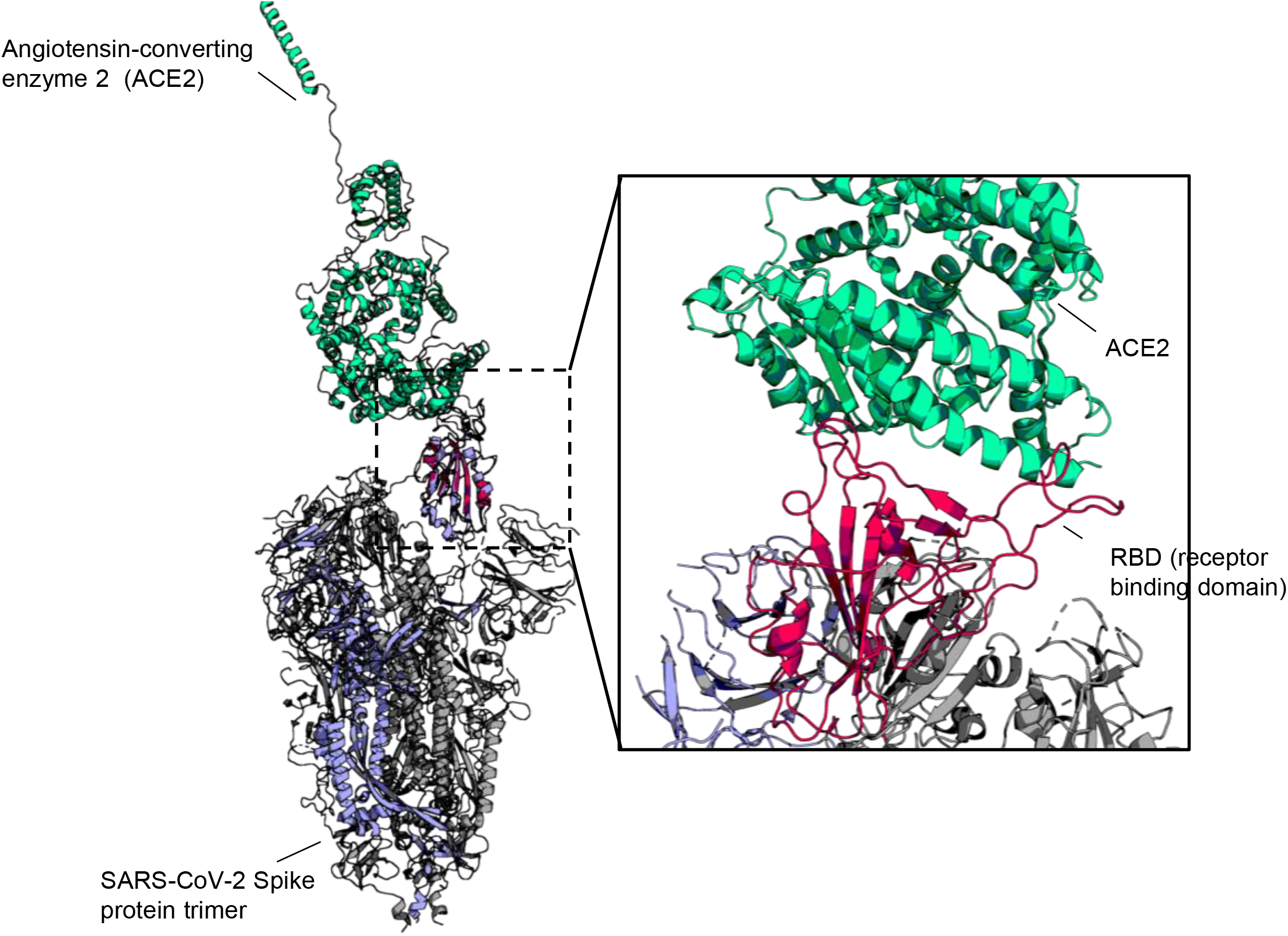
Co-structure of SARS-CoV-2 spike protein RBD with human ACE2. The SARS-CoV-2 spike protein trimer (Marine blue for chain with up-conformation RBD and grey with down-conformation RBD) (PDB code: 6VSB) is aligned to the complex of RBD (red) and human ACE2 (Light green) (PDB code: 6M0J) at the up-conformation RBD to display the binding interface.

**Figure S2b.**
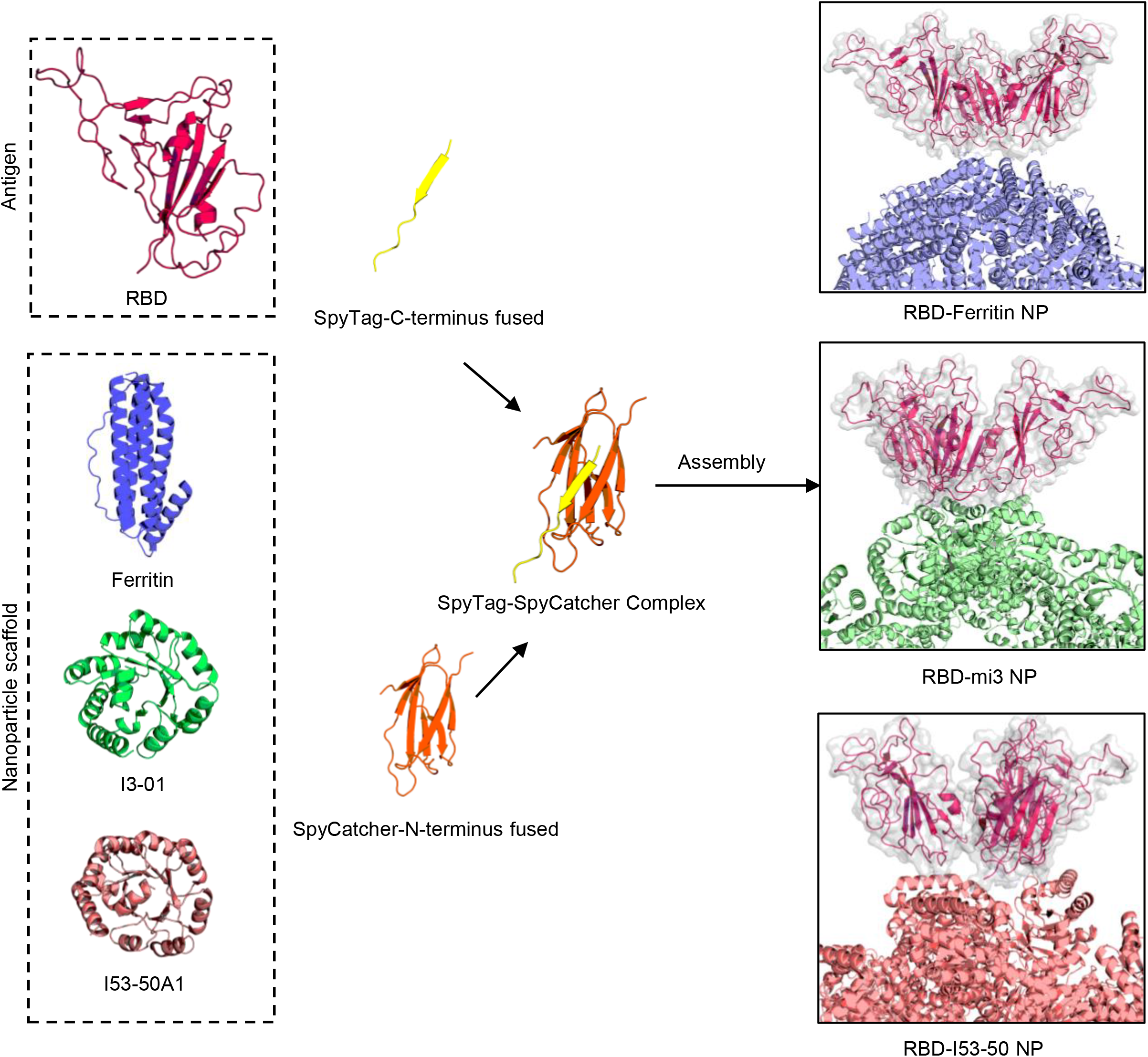
Schematic presentation of covalent bond linking strategy used in RBD-conjugated nanoparticle construction. The RBD fused with SpyTag would automatically links to the ΔN1-SpyCatcher-fused nanoparticle scaffolds ferritin (PDB code: 3BVE), mi3 ^17, 24^ and I53-50 (PDB code: 6P6F) to form a complex of SpyTag-SpyCatcher (PDB code: 4MLI) in between as the bridge. The linked nanoparticles would present RBD (red cartoon with grey surface) on the surface as shown by alignment of C-terminus of RBD with the N-terminus of scaffolds.

**Figure S3.**
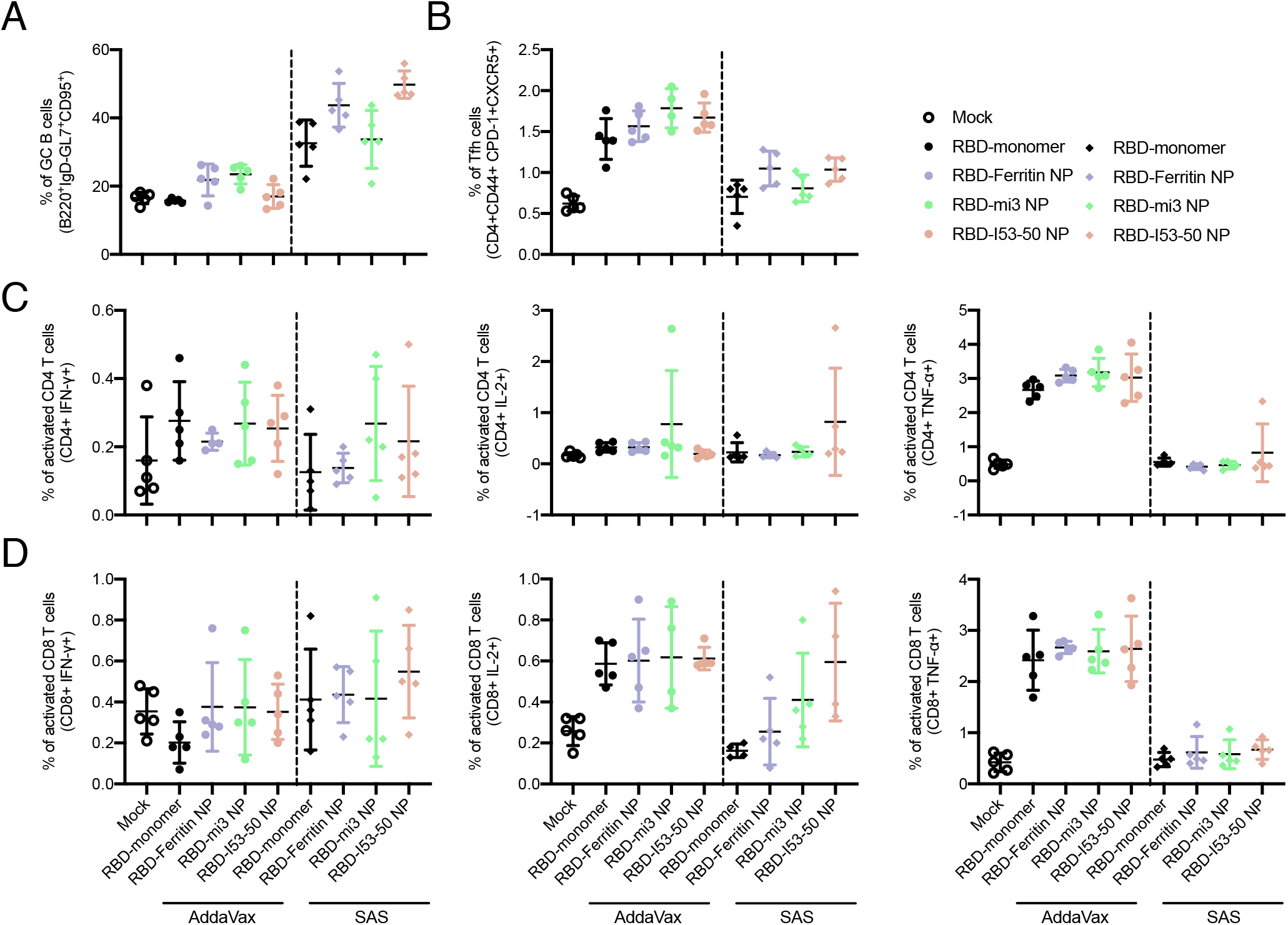
Flowcytometry assay of immune spectrum of drained lymph nodes of immunized mice. (A) Germinal center B cells are marked out from the drained lymph node using B220* (CD45R), IgD-low, GL7* and CD95* as cell marker. Ratio of the positive cells are presented. (B) T follicular helper (Tfh) cells are marked out using CD4*, CD44*, PD-1* and CXCR5* as cell marker. Ratio of the positive cells are presented. (C) (D) Cytokine-secreting CD4* and CD8* T cells are marked out using CD4* and IFN-γ*/IL-2*/TNF-α* as cell markers.

**Figure S4.**
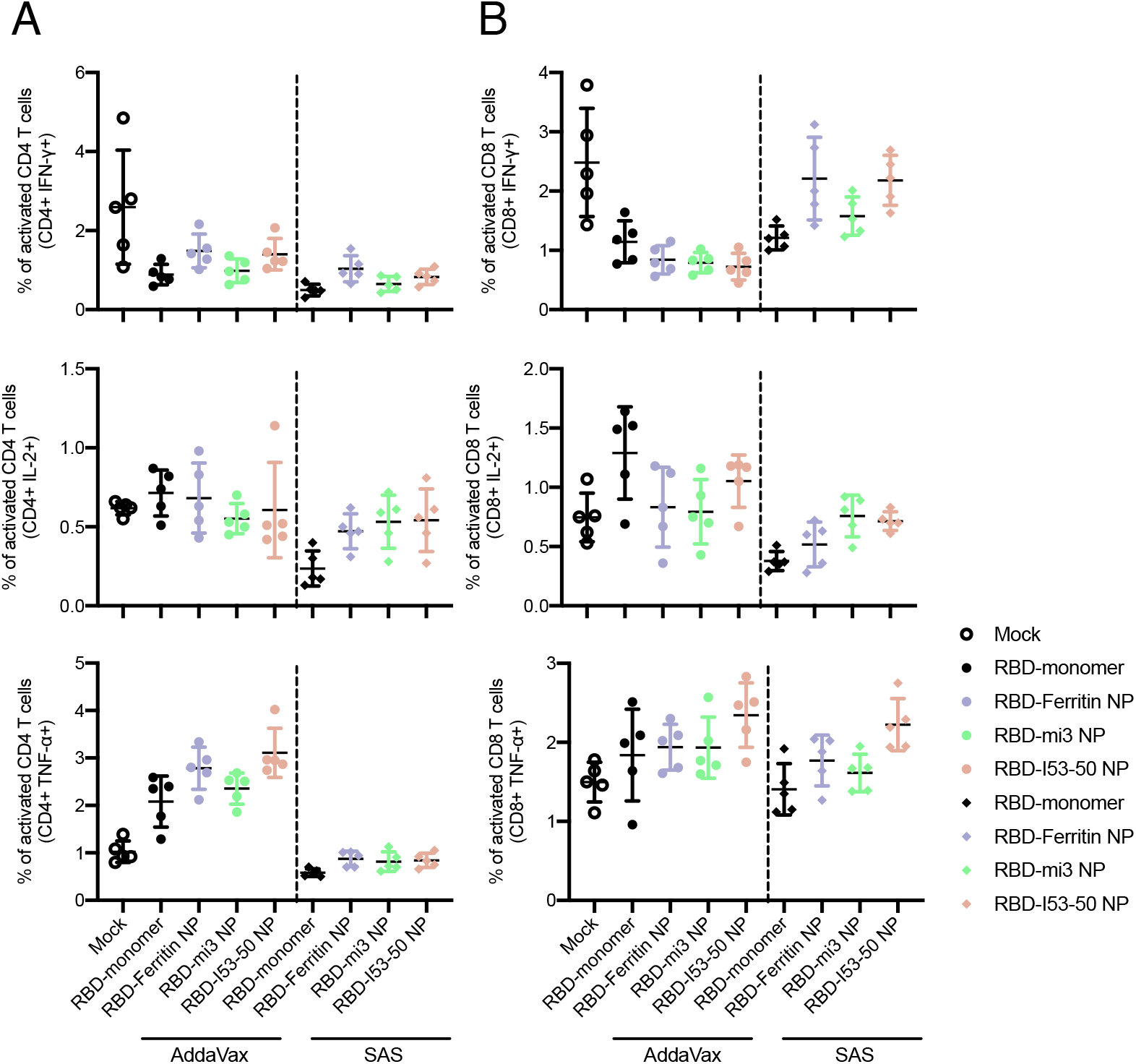
Flowcytometry assay of immune spectrum of spleen of immunized mice. (A)(B) Cytokine-secreting CD4* and CD8* T cells are marked out using CD4* and IFN-γ*/IL-2*/TNF-α* as cell markers.

## Notes

### Competing Interest Statement

The authors have declared no competing interest.

## References

1. de Wit, E.; van Doremalen, N.; Falzarano, D.; Munster, V. J., SARS and MERS: recent insights into emerging coronaviruses. Nat Rev Microbiol 2016, 14 (8), 523–34.

2. Coronaviridae Study Group of the International Committee on Taxonomy of, V., The species Severe acute respiratory syndrome-related coronavirus: classifying 2019-nCoV and naming it SARS-CoV-2. Nat Microbiol 2020, 5 (4), 536–544.

3. Petrosillo, N.; Viceconte, G.; Ergonul, O.; Ippolito, G.; Petersen, E., COVID-19, SARS and MERS: are they closely related? Clin Microbiol Infect 2020, 26 (6), 729–734.

4. Chan, J. F.; Yuan, S.; Kok, K. H.; To, K. K.; Chu, H.; Yang, J.; Xing, F.; Liu, J.; Yip, C. C.; Poon, R. W.; Tsoi, H. W.; Lo, S. K.; Chan, K. H.; Poon, V. K.; Chan, W. M.; Ip, J. D.; Cai, J. P.; Cheng, V. C.; Chen, H.; Hui, C. K.; Yuen, K. Y., A familial cluster of pneumonia associated with the 2019 novel coronavirus indicating person-to-person transmission: a study of a family cluster. Lancet 2020, 395 (10223), 514–523.

5. Wu, F.; Zhao, S.; Yu, B.; Chen, Y. M.; Wang, W.; Song, Z. G.; Hu, Y.; Tao, Z. W.; Tian, J. H.; Pei, Y. Y.; Yuan, M. L.; Zhang, Y. L.; Dai, F. H.; Liu, Y.; Wang, Q. M.; Zheng, J. J.; Xu, L.; Holmes, E. C.; Zhang, Y. Z., A new coronavirus associated with human respiratory disease in China. Nature 2020, 579 (7798), 265–269.

6. Zhou, P.; Yang, X. L.; Wang, X. G.; Hu, B.; Zhang, L.; Zhang, W.; Si, H. R.; Zhu, Y.; Li, B.; Huang, C. L.; Chen, H. D.; Chen, J.; Luo, Y.; Guo, H.; Jiang, R. D.; Liu, M. Q.; Chen, Y.; Shen, X. R.; Wang, X.; Zheng, X. S.; Zhao, K.; Chen, Q. J.; Deng, F.; Liu, L. L.; Yan, B.; Zhan, F. X.; Wang, Y. Y.; Xiao, G. F.; Shi, Z. L., A pneumonia outbreak associated with a new coronavirus of probable bat origin. Nature 2020, 579 (7798), 270–273.

7. Walls, A. C.; Park, Y. J.; Tortorici, M. A.; Wall, A.; McGuire, A. T.; Veesler, D., Structure, Function, and Antigenicity of the SARS-CoV-2 Spike Glycoprotein. Cell 2020, 181 (2), 281–292 e6.

8. Wrapp, D.; Wang, N.; Corbett, K. S.; Goldsmith, J. A.; Hsieh, C. L.; Abiona, O.; Graham, B. S.; McLellan, J. S., Cryo-EM structure of the 2019-nCoV spike in the prefusion conformation. Science 2020, 367 (6483), 1260–1263.

9. Hoffmann, M.; Kleine-Weber, H.; Schroeder, S.; Kruger, N.; Herrler, T.; Erichsen, S.; Schiergens, T. S.; Herrler, G.; Wu, N. H.; Nitsche, A.; Muller, M. A.; Drosten, C.; Pohlmann, S., SARS-CoV-2 Cell Entry Depends on ACE2 and TMPRSS2 and Is Blocked by a Clinically Proven Protease Inhibitor. Cell 2020, 181 (2), 271–280 e8.

10. Li, F.; Li, W.; Farzan, M.; Harrison, S. C., Structure of SARS coronavirus spike receptor-binding domain complexed with receptor. Science 2005, 309 (5742), 1864–8.

11. Lan, J.; Ge, J.; Yu, J.; Shan, S.; Zhou, H.; Fan, S.; Zhang, Q.; Shi, X.; Wang, Q.; Zhang, L.; Wang, X., Structure of the SARS-CoV-2 spike receptor-binding domain bound to the ACE2 receptor. Nature 2020, 581 (7807), 215–220.

12. Shang, J.; Ye, G.; Shi, K.; Wan, Y.; Luo, C.; Aihara, H.; Geng, Q.; Auerbach, A.; Li, F., Structural basis of receptor recognition by SARS-CoV-2. Nature 2020, 581 (7807), 221–224.

13. Wang, Q.; Zhang, Y.; Wu, L.; Niu, S.; Song, C.; Zhang, Z.; Lu, G.; Qiao, C.; Hu, Y.; Yuen, K. Y.; Wang, Q.; Zhou, H.; Yan, J.; Qi, J., Structural and Functional Basis of SARS-CoV-2 Entry by Using Human ACE2. Cell 2020, 181 (4), 894–904 e9.

14. Wang, N.; Shang, J.; Jiang, S.; Du, L., Subunit Vaccines Against Emerging Pathogenic Human Coronaviruses. Front Microbiol 2020, 11, 298.

15. Walls, A. C.; Fiala, B.; Schafer, A.; Wrenn, S.; Pham, M. N.; Murphy, M.; Tse, L. V.; Shehata, L.; O’Connor, M. A.; Chen, C.; Navarro, M. J.; Miranda, M. C.; Pettie, D.; Ravichandran, R.; Kraft, J. C.; Ogohara, C.; Palser, A.; Chalk, S.; Lee, E. C.; Kepl, E.; Chow, C. M.; Sydeman, C.; Hodge, E. A.; Brown, B.; Fuller, J. T.; Dinnon, K. H.; Gralinski, L. E.; Leist, S. R.; Gully, K. L.; Lewis, T. B.; Guttman, M.; Chu, H. Y.; Lee, K. K.; Fuller, D. H.; Baric, R. S.; Kellam, P.; Carter, L.; Pepper, M.; Sheahan, T. P.; Veesler, D.; King, N. P., Elicitation of potent neutralizing antibody responses by designed protein nanoparticle vaccines for SARS-CoV-2. bioRxiv 2020.

16. Dai, L.; Zheng, T.; Xu, K.; Han, Y.; Xu, L.; Huang, E.; An, Y.; Cheng, Y.; Li, S.; Liu, M.; Yang, M.; Li, Y.; Cheng, H.; Yuan, Y.; Zhang, W.; Ke, C.; Wong, G.; Qi, J.; Qin, C.; Yan, J.; Gao, G. F., A Universal Design of Betacoronavirus Vaccines against COVID-19, MERS, and SARS. Cell 2020, 182 (3), 722–733 e11.

17. Bruun, T. U. J.; Andersson, A. C.; Draper, S. J.; Howarth, M., Engineering a Rugged Nanoscaffold To Enhance Plug-and-Display Vaccination. ACS Nano 2018, 12 (9), 8855–8866.

18. Keeble, A. H.; Turkki, P.; Stokes, S.; Khairil Anuar, I. N. A.; Rahikainen, R.; Hytonen, V. P.; Howarth, M., Approaching infinite affinity through engineering of peptide-protein interaction. Proc Natl Acad Sci U S A 2019.

19. Banerjee, A.; Howarth, M., Nanoteamwork: covalent protein assembly beyond duets towards protein ensembles and orchestras. Curr Opin Biotechnol 2018, 51, 16–23.

20. Wang, W.; Zhou, X.; Bian, Y.; Wang, S.; Chai, Q.; Guo, Z.; Wang, Z.; Zhu, P.; Peng, H.; Yan, X.; Li, W.; Fu, Y. X.; Zhu, M., Dual-targeting nanoparticle vaccine elicits a therapeutic antibody response against chronic hepatitis B. Nat Nanotechnol 2020, 15 (5), 406–416.

21. Escolano, A.; Gristick, H. B.; Abernathy, M. E.; Merkenschlager, J.; Gautam, R.; Oliveira, T. Y.; Pai, J.; West, A. P. Jr.; Barnes, C. O.; Cohen, A. A.; Wang, H.; Golijanin, J.; Yost, D.; Keeffe, J. R.; Wang, Z.; Zhao, P.; Yao, K. H.; Bauer, J.; Nogueira, L.; Gao, H.; Voll, A. V.; Montefiori, D. C.; Seaman, M. S.; Gazumyan, A.; Silva, M.; McGuire, A. T.; Stamatatos, L.; Irvine, D. J.; Wells, L.; Martin, M. A.; Bjorkman, P. J.; Nussenzweig, M. C., Immunization expands B cells specific to HIV-1 V3 glycan in mice and macaques. Nature 2019, 570 (7762), 468–473.

22. Yang, J.; Wang, W.; Chen, Z.; Lu, S.; Yang, F.; Bi, Z.; Bao, L.; Mo, F.; Li, X.; Huang, Y.; Hong, W.; Yang, Y.; Zhao, Y.; Ye, F.; Lin, S.; Deng, W.; Chen, H.; Lei, H.; Zhang, Z.; Luo, M.; Gao, H.; Zheng, Y.; Gong, Y.; Jiang, X.; Xu, Y.; Lv, Q.; Li, D.; Wang, M.; Li, F.; Wang, S.; Wang, G.; Yu, P.; Qu, Y.; Yang, L.; Deng, H.; Tong, A.; Li, J.; Wang, Z.; Yang, J.; Shen, G.; Zhao, Z.; Li, Y.; Luo, J.; Liu, H.; Yu, W.; Yang, M.; Xu, J.; Wang, J.; Li, H.; Wang, H.; Kuang, D.; Lin, P.; Hu, Z.; Guo, W.; Cheng, W.; He, Y.; Song, X.; Chen, C.; Xue, Z.; Yao, S.; Chen, L.; Ma, X.; Chen, S.; Gou, M.; Huang, W.; Wang, Y.; Fan, C.; Tian, Z.; Shi, M.; Wang, F. S.; Dai, L.; Wu, M.; Li, G.; Wang, G.; Peng, Y.; Qian, Z.; Huang, C.; Lau, J. Y.; Yang, Z.; Wei, Y.; Cen, X.; Peng, X.; Qin, C.; Zhang, K.; Lu, G.; Wei, X., A vaccine targeting the RBD of the S protein of SARS-CoV-2 induces protective immunity. Nature 2020.

23. Kanekiyo, M.; Bu, W.; Joyce, M. G.; Meng, G.; Whittle, J. R.; Baxa, U.; Yamamoto, T.;Narpala, S.; Todd, J. P.; Rao, S. S.; McDermott, A. B.; Koup, R. A.; Rossmann, M. G.; Mascola, J. R.; Graham, B. S.; Cohen, J. I.; Nabel, G. J., Rational Design of an Epstein-Barr Virus Vaccine Targeting the Receptor-Binding Site. Cell 2015, 162 (5), 1090–100.

24. Hsia, Y.; Bale, J. B.; Gonen, S.; Shi, D.; Sheffler, W.; Fong, K. K.; Nattermann, U.; Xu, C.; Huang, P. S.; Ravichandran, R.; Yi, S.; Davis, T. N.; Gonen, T.; King, N. P.; Baker, D., Corrigendum: Design of a hyperstable 60-subunit protein icosahedron. Nature 2016, 540 (7631), 150.

25. Bale, J. B.; Gonen, S.; Liu, Y.; Sheffler, W.; Ellis, D.; Thomas, C.; Cascio, D.; Yeates, T. O.; Gonen, T.; King, N. P.; Baker, D., Accurate design of megadalton-scale two-component icosahedral protein complexes. Science 2016, 353 (6297), 389–94.

26. Du, L.; Zhao, G.; Chan, C. C.; Sun, S.; Chen, M.; Liu, Z.; Guo, H.; He, Y.; Zhou, Y.; Zheng, B. J.; Jiang, S., Recombinant receptor-binding domain of SARS-CoV spike protein expressed in mammalian, insect and E.coli cells elicits potent neutralizing antibody and protective immunity. Virology 2009, 393 (1), 144–50.

27. Shi, R.; Shan, C.; Duan, X.; Chen, Z.; Liu, P.; Song, J.; Song, T.; Bi, X.; Han, C.; Wu, L.;Gao, G.; Hu, X.; Zhang, Y.; Tong, Z.; Huang, W.; Liu, W. J.; Wu, G.; Zhang, B.; Wang, L.; Qi, J.; Feng, H.; Wang, F. S.; Wang, Q.; Gao, G. F.; Yuan, Z.; Yan, J., A human neutralizing antibody targets the receptor-binding site of SARS-CoV-2. Nature 2020, 584 (7819), 120–124.

28. Mills, C. D.; Kincaid, K.; Alt, J. M.; Heilman, M. J.; Hill, A. M., M-1/M-2 macrophages and the Th1/Th2 paradigm. J Immunol 2000, 164 (12), 6166–73.

29. Amanat, F.; Krammer, F., SARS-CoV-2 Vaccines: Status Report. Immunity 2020, 52 (4), 583–589.

30. Edwards, K. M., Vaccines targeting SARS-CoV-2 tested in humans. Nat Med 2020, 26 (9), 1336–1338.

31. Graham, B. S., Rapid COVID-19 vaccine development. Science 2020, 368 (6494), 945–946.

32. Ng, W. H.; Liu, X.; Mahalingam, S., Development of vaccines for SARS-CoV-2. F1000Res 2020, 9.

33. Hou, Y. J.; Okuda, K.; Edwards, C. E.; Martinez, D. R.; Asakura, T.; Dinnon, K. H. 3rd; Kato, T.; Lee, R. E.; Yount, B. L.; Mascenik, T. M.; Chen, G.; Olivier, K. N.; Ghio, A.; Tse, L. V.; Leist, S. R.; Gralinski, L. E.; Schafer, A.; Dang, H.; Gilmore, R.; Nakano, S.; Sun, L.; Fulcher, M. L.; Livraghi-Butrico, A.; Nicely, N. I.; Cameron, M.; Cameron, C.; Kelvin, D. J.; de Silva, A.; Margolis, D. M.; Markmann, A.; Bartelt, L.; Zumwalt, R.; Martinez, F. J.; Salvatore, S. P.; Borczuk, A.; Tata, P. R.; Sontake, V.; Kimple, A.; Jaspers, I.; O’Neal, W. K.; Randell, S. H.; Boucher, R. C.; Baric, R. S., SARS-CoV-2 Reverse Genetics Reveals a Variable Infection Gradient in the Respiratory Tract. Cell 2020, 182 (2), 429–446 e14.

34. Thi Nhu Thao, T.; Labroussaa, F.; Ebert, N.; V’Kovski, P.; Stalder, H.; Portmann, J.; Kelly, J.; Steiner, S.; Holwerda, M.; Kratzel, A.; Gultom, M.; Schmied, K.; Laloli, L.; Husser, L.; Wider, M.; Pfaender, S.; Hirt, D.; Cippa, V.; Crespo-Pomar, S.; Schroder, S.; Muth, D.; Niemeyer, D.; Corman, V. M.; Muller, M. A.; Drosten, C.; Dijkman, R.; Jores, J.; Thiel, V., Rapid reconstruction of SARS-CoV-2 using a synthetic genomics platform. Nature 2020, 582 (7813), 561–565.

35. Case, J. B.; Rothlauf, P. W.; Chen, R. E.; Liu, Z.; Zhao, H.; Kim, A. S.; Bloyet, L. M.; Zeng, Q.; Tahan, S.; Droit, L.; Ilagan, M. X. G.; Tartell, M. A.; Amarasinghe, G.; Henderson, J. P.; Miersch, S.; Ustav, M.; Sidhu, S.; Virgin, H. W.; Wang, D.; Ding, S.; Corti, D.; Theel, E. S.; Fremont, D. H.; Diamond, M. S.; Whelan, S. P. J., Neutralizing Antibody and Soluble ACE2 Inhibition of a Replication-Competent VSV-SARS-CoV-2 and a Clinical Isolate of SARS-CoV-2. Cell Host Microbe 2020, 28 (3), 475–485 e5.

36. Mercado, N. B.; Zahn, R.; Wegmann, F.; Loos, C.; Chandrashekar, A.; Yu, J.; Liu, J.; Peter, L.; McMahan, K.; Tostanoski, L. H.; He, X.; Martinez, D. R.; Rutten, L.; Bos, R.; van Manen, D.; Vellinga, J.; Custers, J.; Langedijk, J. P.; Kwaks, T.; Bakkers, M. J.G.; Zuijdgeest, D.; Rosendahl Huber, S. K.; Atyeo, C.; Fischinger, S.; Burke, J. S.; Feldman, J.; Hauser, B. M.; Caradonna, T. M.; Bondzie, E. A.; Dagotto, G.; Gebre, M. S.; Hoffman, E.; Jacob-Dolan, C.; Kirilova, M.; Li, Z.; Lin, Z.; Mahrokhian, S. H.; Maxfield, L. F.; Nampanya, F.; Nityanandam, R.; Nkolola, J. P.; Patel, S.; Ventura, J. D.; Verrington, K.; Wan, H.; Pessaint, L.; Van Ry, A.; Blade, K.; Strasbaugh, A.; Cabus, M.; Brown, R.; Cook, A.; Zouantchangadou, S.; Teow, E.; Andersen, H.; Lewis, M. G.; Cai, Y.; Chen, B.; Schmidt, A. G.; Reeves, R. K.; Baric, R. S.; Lauffenburger, D. A.; Alter, G.; Stoffels, P.; Mammen, M.; Van Hoof, J.; Schuitemaker, H.; Barouch, D. H., Single-shot Ad26 vaccine protects against SARS-CoV-2 in rhesus macaques. Nature 2020.

37. Zhang, N. N.; Li, X. F.; Deng, Y. Q.; Zhao, H.; Huang, Y. J.; Yang, G.; Huang, W. J.; Gao, P.; Zhou, C.; Zhang, R. R.; Guo, Y.; Sun, S. H.; Fan, H.; Zu, S. L.; Chen, Q.; He, Q.; Cao, T. S.; Huang, X. Y.; Qiu, H. Y.; Nie, J. H.; Jiang, Y.; Yan, H. Y.; Ye, Q.; Zhong, X.; Xue, X. L.; Zha, Z. Y.; Zhou, D.; Yang, X.; Wang, Y. C.; Ying, B.; Qin, C. F., A Thermostable mRNA Vaccine against COVID-19. Cell 2020, 182 (5), 1271–1283 e16.

38. Gao, Q.; Bao, L.; Mao, H.; Wang, L.; Xu, K.; Yang, M.; Li, Y.; Zhu, L.; Wang, N.; Lv, Z.; Gao, H.; Ge, X.; Kan, B.; Hu, Y.; Liu, J.; Cai, F.; Jiang, D.; Yin, Y.; Qin, C.; Li, J.; Gong, X.; Lou, X.; Shi, W.; Wu, D.; Zhang, H.; Zhu, L.; Deng, W.; Li, Y.; Lu, J.; Li, C.; Wang, X.; Yin, W.; Zhang, Y.; Qin, C., Development of an inactivated vaccine candidate for SARS-CoV-2. Science 2020, 369 (6499), 77–81.

39. Wang, H.; Zhang, Y.; Huang, B.; Deng, W.; Quan, Y.; Wang, W.; Xu, W.; Zhao, Y.; Li, N.; Zhang, J.; Liang, H.; Bao, L.; Xu, Y.; Ding, L.; Zhou, W.; Gao, H.; Liu, J.; Niu, P.; Zhao, L.; Zhen, W.; Fu, H.; Yu, S.; Zhang, Z.; Xu, G.; Li, C.; Lou, Z.; Xu, M.; Qin, C.; Wu, G.; Gao, G. F.; Tan, W.; Yang, X., Development of an Inactivated Vaccine Candidate, BBIBP-CorV, with Potent Protection against SARS-CoV-2. Cell 2020, 182 (3), 713–721 e9.

40. Rappuoli, R.; Serruto, D., Self-Assembling Nanoparticles Usher in a New Era of Vaccine Design. Cell 2019, 176 (6), 1245–1247.

41. Irvine, D. J.; Hanson, M. C.; Rakhra, K.; Tokatlian, T., Synthetic Nanoparticles for Vaccines and Immunotherapy. Chem Rev 2015, 115 (19), 11109–46.

42. Huang, P. S.; Boyken, S. E.; Baker, D., The coming of age of de novo protein design. Nature 2016, 537 (7620), 320–7.

43. Du, L.; He, Y.; Zhou, Y.; Liu, S.; Zheng, B. J.; Jiang, S., The spike protein of SARS-CoV--a target for vaccine and therapeutic development. Nat Rev Microbiol 2009, 7 (3), 226–36.

44. Starr, T. N.; Greaney, A. J.; Hilton, S. K.; Ellis, D.; Crawford, K. H. D.; Dingens, A. S.; Navarro, M. J.; Bowen, J. E.; Tortorici, M. A.; Walls, A. C.; King, N. P.; Veesler, D.; Bloom, J. D., Deep Mutational Scanning of SARS-CoV-2 Receptor Binding Domain Reveals Constraints on Folding and ACE2 Binding. Cell 2020, 182 (5), 1295–1310 e20.

45. Arvin, A. M.; Fink, K.; Schmid, M. A.; Cathcart, A.; Spreafico, R.; Havenar-Daughton, C.; Lanzavecchia, A.; Corti, D.; Virgin, H. W., A perspective on potential antibody-dependent enhancement of SARS-CoV-2. Nature 2020, 584 (7821), 353–363.

46. Wang, Y.; Wang, L.; Cao, H.; Liu, C., SARS-CoV-2 S1 is superior to the RBD as a COVID-19 subunit vaccine antigen. J Med Virol 2020.

47. Bachmann, M. F.; Jennings, G. T., Vaccine delivery: a matter of size, geometry, kinetics and molecular patterns. Nat Rev Immunol 2010, 10 (11), 787–96.

48. Graham, B. S.; Gilman, M. S.A.; McLellan, J. S., Structure-Based Vaccine Antigen Design. Annu Rev Med 2019, 70, 91–104.

49. Zhang, X.; Zhao, B.; Ding, M.; Song, S.; Kang, Y.; Yu, Y.; Xu, M.; Xiang, T.; Gao, L.; Feng, Q.; Zhao, Q.; Zeng, M. S.; Krummenacher, C.; Zeng, Y. X., A novel vaccine candidate based on chimeric virus-like particle displaying multiple conserved epitope peptides induced neutralizing antibodies against EBV infection. Theranostics 2020, 10 (13), 5704–5718.

50. Ou, X.; Liu, Y.; Lei, X.; Li, P.; Mi, D.; Ren, L.; Guo, L.; Guo, R.; Chen, T.; Hu, J.; Xiang, Z.; Mu, Z.; Chen, X.; Chen, J.; Hu, K.; Jin, Q.; Wang, J.; Qian, Z., Characterization of spike glycoprotein of SARS-CoV-2 on virus entry and its immune cross-reactivity with SARS-CoV. Nat Commun 2020, 11 (1), 1620.

51. Robert, X.; Gouet, P., Deciphering key features in protein structures with the new ENDscript server. Nucleic Acids Res 2014, 42 (Web Server issue), W320–4.

